# Phenotypic plasticity as a route to population shifts via tipping points

**DOI:** 10.64898/2026.04.14.718490

**Authors:** Benedict C. Fellows, Steven M. White, Dominic P. Brass, Alexander Nascou, Christina A. Cobbold

## Abstract

Environmental change has caused dramatic global declines in biodiversity, with some species showing abrupt and often irreversible changes in population abundance. These regime shifts can occur when environmental thresholds, known as tipping points, are passed. Many species can respond to environmental change via phenotypic plasticity with the expectation that strong phenotypic plasticity reduces the risk of regime shifts by enabling a species to rapidly respond to environmental change, potentially mitigating the risks of population collapse. Testing the theory that phenotypic plasticity buffers against regime shifts requires a novel whole population approach that robustly considers the feedback mechanisms between environment, phenotype and population density, common to the life-history of many species. For this purpose we develop a tractable mathematical framework, and demonstrate, counter-intuitively, that phenotypic plasticity can induce tipping points, due to the inclusion of feedback mechanisms that operate at both the level of the organism and population. Consequently, predicting the existence of potentially devastating tipping points and so understanding ecosystem collapse is more nuanced than current thinking suggests.

## 1 Introduction

Our natural environment is under considerable stress driven by a wide range of anthropogenic factors including climate change (Cahill *et al*., 2013; Parmesan *et al*., 2023), loss of habitat (Horgan, 2005; Bojsen and Barriga, 2002; McKinney, 2008), pollution (Scheffer *et al*., 1993), and over-harvesting (Möllmann *et al*., 2021), resulting in an increasing risk of ecosystem collapse. Establishing how resilient ecosystems are to environmental stress is vital to improve environmental monitoring and management strategies (Oliver *et al*., 2015; Biggs *et al*., 2012). However, in many ecosystems, small changes in environmental stressors can lead to abrupt changes in the ecosystem state (regime shifts) (Scheffer and Carpenter, 2003; Turner *et al*., 2020), which are becoming increasingly common (Turner *et al*., 2020; Folke *et al*., 2004; Ratajczak *et al*., 2018). The environmental thresholds triggering these abrupt shifts in ecosystems are known as tipping points (van Nes *et al*., 2016; Dakos *et al*., 2019). Evidence of tipping points is observed across ecosystems, including tropical forest to savanna transitions (Hirota *et al*., 2011; Staal *et al*., 2020; Oliveras and Malhi, 2016), the dieback of boreal forests into steppes (Scheffer *et al*., 2012; Rao *et al*., 2023), shifts to algal dominance (eutrophication) in water bodies (Hughes, 1994; Graham *et al*., 2015; Holbrook *et al*., 2016), and collapses of cod fisheries (Hutchings and Reynolds, 2004; Winter *et al*., 2020; Möllmann *et al*., 2021). Following a collapse, restoring the ecosystem to the original state often requires a change in stressor substantially greater than the change that induced the collapse (Mumby *et al*., 2007). In the worst case ecosystems might not be recoverable (Hutchings and Reynolds, 2004; Petraitis, 2013).

Ecological theory suggests tipping points often arise when there is a collapse of bistability, due to a weakening of the strong positive feedback loops maintaining two alternative stable ecosystem states (see black line in Figure 1a and Scheffer and Carpenter (2003); Clements and Ozgul (2018); May (1977)). Such tipping points typify hysteresis, which differs from smooth critical transitions between ecosystem states (red line in Figure 1b); once a tipping point is crossed, critical transitions and collapse cannot easily be reversed. Dynamical models of tipping points have primarily been developed for systems where the population has fixed traits (Schreiber and Rudolf, 2008; Hao *et al*., 2026). However, many organisms possess traits that allow the population to respond to stress that could buffer the population against critical transitions, by allowing individuals to shift their traits to maintain population fitness. At the population level there is evidence from microcosm experiments (Clements and Ozgul, 2016) and field populations in fisheries (Clements *et al*., 2017) of significant trait change prior to critical population transitions, although the mechanism for the observed trait changes is yet to be established.

**Figure 1:**
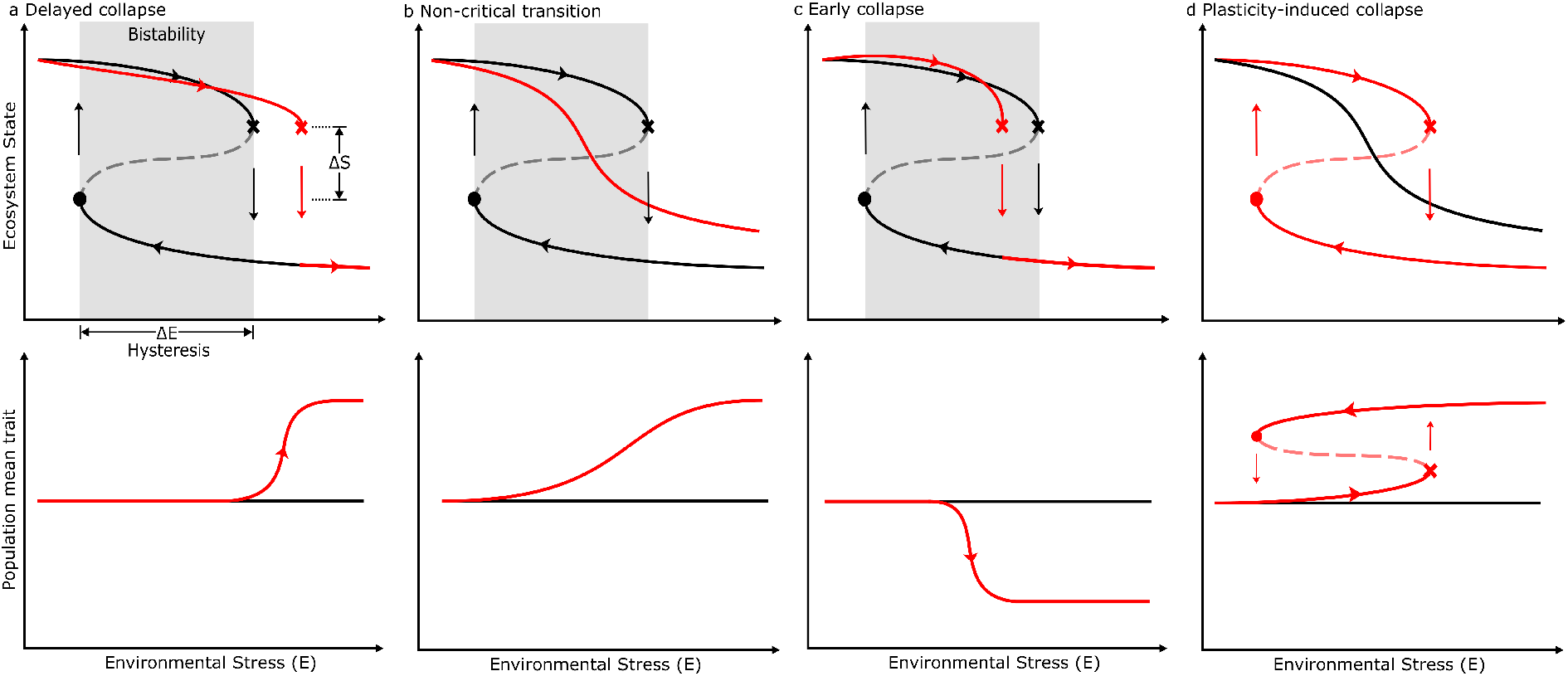
Potential patterns of critical transitions in abundance and plastic traits in response to changing environmental conditions. In all plots tipping points at high and low environmental stress are denoted by crosses and circles respectively, between these values alternative stable states exist (solid lines) and the width of this shaded region (Δ*E*) indicates the magnitude of environmental change required to return the system to it’s prior state. Arrows indicate the transition of the ecosystem state as environmental stress is changed. The magnitude of the change in ecosystem state between the two tipping points is denoted by Δ*S*. Columns (**a**)-(**c**): In the absence of phenotypic plasticity (black lines), critical transitions and discontinuous changes in ecosystem state occur at tipping points in environmental stress, *E*. In the presence of phenotypic plasticity (red lines), trait change may: (**a**), delay the onset of tipping points, (**b**) provide a smooth pathway to non-critical transitions or (**c**) lead to early onset of tipping points. By contrast in (**d**), if in the absence of plasticity the ecosystem does **not** exhibit tipping points (black line), we hypothesise phenotypic plasticity may induce tipping points, creating critical transitions in abundance and trait that were otherwise absent (red line), resulting in hysteresis in both population abundance and trait. The missing sections of the red hysteresis curves in (**a-c**) reflect further gaps in our current knowledge, there is speculation in the literature over whether plasticity would shift one or both tipping points.

Phenotypic plasticity, the ability for individual organisms to express different phenotypes (observable traits) depending on environmental conditions (Pigliucci *et al*., 2006; Fusco and Minelli, 2010; Sommer, 2020), is an example of trait change through which individuals can rapidly respond to changes in the environment. Phenotypic plasticity is often considered to be a means of buffering environmental change, and reducing environmental variance in population growth (Diamond and Martin, 2021). The capacity of phenotypic plasticity to buffer populations in this way is typically described by the degree to which a phenotypic trait can be changed (Burton *et al*., 2022). The characteristics of phenotypic plasticity vary widely from instantaneous reversible plasticity (e.g., temperature dependent flight behaviour in fruit flies (Lehmann, 1999)) to delayed irreversible plasticity (e.g., developmental plasticity where environmental conditions in one life-stage affect traits in later stages (Schneider, 2022)). This rapid mechanism of environmental response is exhibited across all taxa, including mammals (Boutin and Lane, 2014), birds (Charmantier and Gienapp, 2014), fish (Crozier and Hutchings, 2014; Seebacher *et al*., 2015), invertebrates (Seebacher *et al*., 2015; Stoks *et al*., 2014), protists (Gibert *et al*., 2022) and plants (Franks *et al*., 2014). It has been proposed that phenotypic plasticity could delay the onset of tipping points, shifting the threshold for critical transitions to a higher level of environmental stress or allow for a smooth (and reversible) transition between ecosystem states (see the red line in Figure 1a and b and Cerini *et al*. (2023); Dakos *et al*. (2019); Carvalho *et al*. (2023)). However, when the relationship between environmental stress and trait plasticity is non-linear, phenotypic responses are constrained and the ability of phenotypic plasticity to mitigate the effects of environmental stressors is limited (Cerini *et al*., 2023; Clements and Ozgul, 2018). If environment induced trait changes have a negative impact on the average fitness of the population, plasticity may lower the threshold level of environmental stress that triggers a critical transition, leading to an earlier onset of tipping points (see the red line in Figure 1c and Dakos *et al*. (2019)). Phenotypic plasticity is not only a response to abiotic stresses on a population, but can also be induced by biotic environmental stresses linked to population density (Foquet *et al*., 2021). Juvenile competition for resources can lead to plasticity in the form of size dependent survival and reproduction in later life-stages (Couret *et al*., 2014), with high performance of individuals emerging from higher food or lower density juvenile environments (van Allen *et al*., 2010). Such density-dependent trait responses are embedded in complex life cycles and provide a mechanism for density-dependent feedbacks that can lead to counterintuitive population responses to environmental change (de Roos, 2020; Edwards and Smallegange, 2025). Here, we postulate that if the expression of a phenotype is dependent on density, a strong positive feedback between trait and population abundance could induce tipping points and regime shifts, which would not otherwise be present (see the red line in Figure 1d). Moreover, we hypothesise that a high degree of plasticity (a strong interaction between trait and environment and a high capacity for trait change), may increase the strength of density-dependent feedbacks and strengthen hysteresis rather than buffer the population from environmental change.

To test our hypotheses we use a robust tractable framework for modelling populations that allows for irreversible developmental plasticity (by using a system of stage-structured delay-differential equations) to explore the role that phenotypic plasticity has on the creation of tipping points. Our approach links experimentally derived environment–trait relationships to a well-parametrised and empirically motivated stage-structured model. Via both analytic and numerical approaches and a set of simplified models, we isolate the effects of phenotypic plasticity from other mechanisms that are known to induce tipping points (e.g., ontogenetic niche shifts (Schreiber and Rudolf, 2008)) and demonstrate a role for phenotypic plasticity in generating tipping points and population collapse. By varying the shape of the environment-trait relationships (reaction norms), that capture the sensitivity of traits to changes in the environment, we find that the capacity for trait variation in the population is what maintains an alternative stable system state, highlighting the importance of individual life-history and environment-trait relationships on our understanding of such regime shifts.

## 2 Material and methods

As an exemplar system for studying the influence of phenotypic plasticity in generating population tipping points we consider Nicholson’s blowfly (*Lucilia cuprina*) experiments (Nicholson, 1957) in which, counter-intuitively, increasing food caused a regime shift and reduced the population abundance by 50% (Nicholson, 1957). Nicholson made heuristic arguments for a role for phenotypic plasticity in generating these complex population dynamics. The phenotypic plasticity that he referred to is in response to larval food availability, and affects survival and fecundity life-history traits. This presents as delayed irreversible phenotypic plasticity; larvae that acquire more food are larger, are better able to survive through pupation, and become larger adults with higher maximum adult fecundity. In Nicholson’s experiments blowfly experience competition for food in both the larval and adult stages, although they consume different resources (they undergo an ontogenetic niche shift), and limitation of adult food can result in fewer eggs being produced, reducing competition in the larval stage so that larvae acquire more food, creating a feedback loop as argued by Nicholson. The combination of delayed developmental plasticity and ontogenetic niche shift are common to many species, making the blowfly system a good model system to explore the mechanism of plasticity induced regime shifts. Ontogenetic niche shifts are known to cause hysteresis leading to tipping points (Schreiber and Rudolf, 2008) and so to isolate the effects of plasticity from those of the niche-shift, in addition to the main model of blowfly dynamics, we also consider a number of simplified models that enable us to turn off plasticity and the ontogenetic niche shift in turn. A further simplified model allows us to explore the effects of the degree of plasticity by changing the sensitivity of traits to changes in the environment.

### 2.1 Main model

Many models have been developed to understand the ecological drivers of Nicholson’s experimental findings (Gurney *et al*., 1983; May, 1986; Glyzin, 2017). We adopt the model of Brass *et al*. (2021) which explicitly includes phenotypic plasticity and reproduces Nicholson’s experimental observations. The mathematical model is a system of stage and phenotype-structured delay-differential equations, where cohorts of individuals with a shared phenotype are tracked through development, incorporating the life-stage durations via time delays. Within this model *Lucilia cuprina* is assumed to have 5 main life-stages: eggs, larvae, pupae, non-reproductive juvenile adults and reproductive adults. Eggs, pupae and non-reproductive adults can be modelled implicitly as these stages only experience density-independent mortality and therefore simply act as time delays for development, with a fixed proportion surviving each of these developmental stages, consequently the problem is reduced to modelling two life-stages (depicted in Figure 2): the larval stage (*L*) and the reproductive adult stage (*A*_*i*_) (referred to as adults). The adults are further partitioned into *n* size-classes determined by the food consumed during larval development.

**Figure 2:**
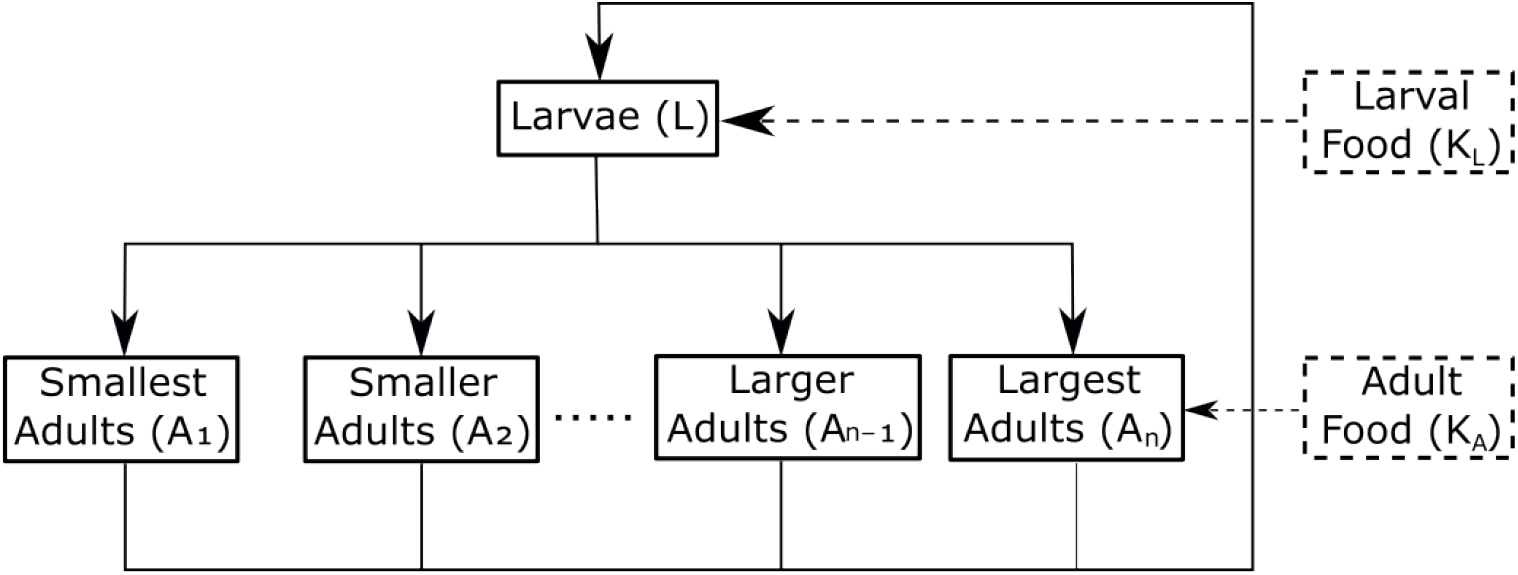
Model schematic. The population is partitioned into two stages: Larvae (*L*) and a size-discretised reproductive adult stage (*A*_*i*_), ranging from the smallest adult size class *A*_1_ to the largest adult size class *A*_*n*_. The egg, pupal and non-reproductive adult stages are modelled implicitly. There are two food sources in the model: the daily larval food supply (*K*_*L*_) and daily adult food supply (*K*_*A*_). Average larval food per capita obtained over the duration of the larval stage determines the adult size and the traits of through-pupal survival and maximum adult fecundity.

Adults in size class *i* are assumed to have a shared phenotype based on the environmental stress experienced during their larval development, where more larval food per capita leads to an increased adult size. The phenotype of individuals in adult size class *i* is composed of the two life-history traits: through-pupal survival proportion *S*_*Pi*_, which describes survival for an individual from pupation until maturation into a non-reproductive adult, and maximum adult fecundity *q*_*i*_, which captures the egg laying rate of individual adults in an unlimited food environment. A constant amount of adult and larval food is supplied daily denoted by *K*_*A*_ and *K*_*L*_ respectively, varying food supply acts to change the environmental conditions experienced by the blowfly and to account for the delays in the development between each life-stage, the system is modelled using a set of delay differential equations:

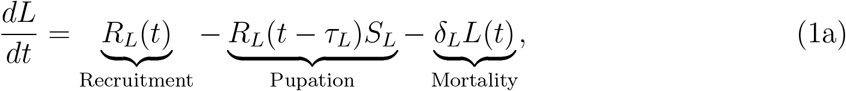

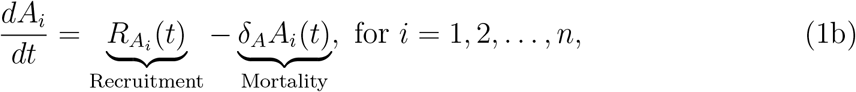

where the recruitment terms for larvae and adults in size class *i* are given by:

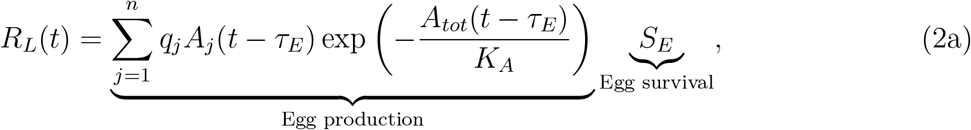

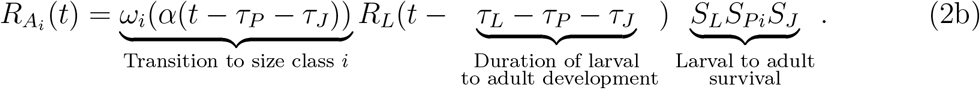

Recruitment into the larval stage is dependent on adult fecundity. An adult fly has a number, *q*_*i*_, of ovarioles based upon its size, but a proportion, 1 *−* exp(*−A*_*tot*_(*t − τ*_*E*_)*/K*_*A*_), under go resorption if the adult does not get enough food to mature them, where *A*_*tot*_ = Σ _*j*_ *A*_*j*_ is the total abundance of adults across all size classes. The function capturing this density-dependent effect is taken from Gurney *et al*. (1983) who fit laboratory data of egg production as a function of adult food supply rate (Figure 2b of Gurney *et al*. (1983)), competition was scramble-like and we assume that reproductive adults compete equally for the available adult food (*K*_*A*_) with a consumption rate of 1 mg per day per adult, regardless of phenotype. A proportion *S*_*E*_ of the eggs laid then survive the egg development period (*τ*_*E*_) to mature into larvae.

Each life-stage prior to reproductive maturity (*I* = *E, L, P, J* for eggs, larvae, pupae and non-reproductive adults respectively) has a corresponding development time *τ*_*I*_. Eggs, larvae, non-reproductive adults (*I* = *E, L, J*), have through-stage survival proportion given by *S*_*I*_ = exp(*−δ*_*I*_*τ*_*I*_), where *δ*_*I*_ is the associated density independent mortality rate and for adults this rate is *δ*_*A*_. Recruitment into the adult stage requires survival from larvae to adult, where the through-pupal survival proportion *S*_*Pi*_ depends on the pupal size determined by the food consumed as a larvae. A constant amount of larval food, *K*_*L*_, is supplied daily and the average larval food per capita over the larval development period is determined by scramble competition for the limited food resource, as such it is correlated to larval density and satisfies

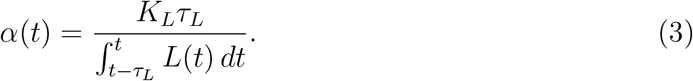

There is no evidence to suggest that larval competition affects larval survival and it is not included in the model. Finally, the transition function, *ω*_*i*_(*α*(*t*)), assigns individuals to adult size class *i* based on the average daily amount of food consumed as a larvae and is given by:

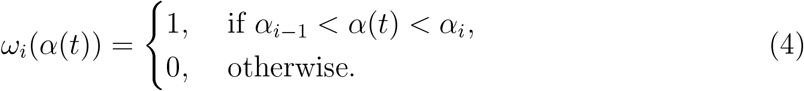

The choice of *ω*_*i*_ precludes individual variation in trait expression, this is to enable the analysis to isolate the role of phenotypic plasticity from other sources of variation e.g., genetic. Higher levels of larval food consumption result in entry into a larger adult size class as observed in Nicholson’s experiments.

The values for the life-history traits for each adult size class *i* are determined by discretising the reaction norms *S*_*P*_ (*α*) and *q*(*α*), that describe the relationship between these traits and (per capita) food acquired during the larval stage (*α*), a measure of environmental stress during early development. Adult size-class *i*, is characterised by a minimum (*α*_*i−*1_) and maximum (*α*_*i*_) average level of larval food obtained per capita required to reach size class *i*. The effect of the average protein available per larva per day on the traits expressed by blowflies is expected to saturate at *α*_*max*_ = 132 mg per larva per day and we expect that at levels below *α*_*min*_ = 1.15 all larvae starve (Brass *et al*., 2021). Therefore, since the maximum adult fecundity, *q*(*α*), is a strictly increasing function of *α* with a minimum (*q*_*min*_ = *q*(*α*_*min*_) and maximum *q*_*max*_ = *q*(*α*_*max*_)) we discretise the adults into *n* size classes such that the difference in *q*(*α*) between consecutive size classes is fixed at (*q*_*max*_ *− q*_*min*_)*/n*, with *n* chosen sufficiently large to discretise the continuous environment-trait reaction norms.

Thus,

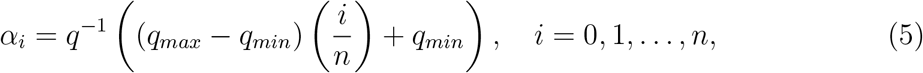

where *q*^*−*1^(*α*) denotes the inverse of *q*(*α*). Each adult size-class, *i*, has associated traits for maximum adult fecundity, *q*_*i*_, and through-pupal survival, *S*_*Pi*_, determined by the midpoints 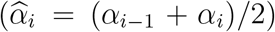 of the intervals (*α*_*i−*1_,*α*_*i*_), specifically 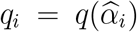and 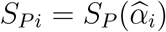

The two non-linear reaction norms, *S*_*P*_ (*α*) and *q*(*α*) are adapted from parametrisation of laboratory data by Moe *et al*. (2002) and Webber (1955) and illustrated in Figure 4f-g. Moe *et al*. (2002) found through-pupal survival to have a delayed effect on larval density; survival was highest at low initial larval densities and declined at higher initial larval densities, with a quadratic effect of density. We take the adapted version of the functional response parametrised using quadratic logistic regression in Moe *et al*. (2002). The adaptation developed by Brass *et al*. (2021) transforms food so that it is quantified as average protein available per larvae per day to be consistent with Nicholson’s experiments, giving

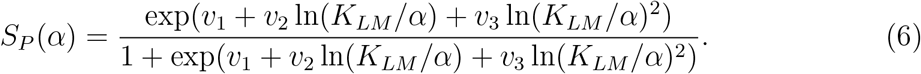

The hump shaped survival function is a result of an Allee effect, where at low densities (high per capita food) larvae have not been able benefit from efficient feeding that comes from being in a larger group, reducing pupal size and pupal survival (Moe *et al*., 2002).

The maximum adult fecundity (*q*(*α*)) also has a delayed effect of larval density, whereby higher larval densities resulted in smaller adults with fewer ovarioles, the relationship parameterised by Webber (1955) is described by

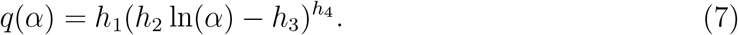

The parameters *v*_*i*_ and *h*_*i*_ control the shape of the two reaction norms, with *v*_3_, *h*_3_ and *h*_4_ controlling the higher order nonlinearity. The fitted values are given in Supplementary Table S.1.1.

### 2.2 Model simplifications

We examine three model simplifications that allow us to (1) determine the effects of the ontogenetic niche shift without phenotypic plasticity, (2) determine the effects of phenotypic plasticity without the ontogenetic niche shift, and (3) determine the effects of the degree of plasticity in through-pupal survival, which we demonstrate is central in determining the existence of tipping points.

#### (1) Fixed trait model: No phenotypic plasticity

In the fixed trait model size dependent effects of larval competition are removed and there is no phenotypic plasticity. We assume all individuals, regardless of food acquired as a larvae, have the same fixed trait values:

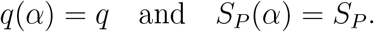

The ontogenetic niche shift remains, but availability of larval resource has no limiting effects on the population.

#### (2) Shared resource model: No ontogenetic niche shift

To isolate effects of phenotypic plasticity we remove the ontogenetic niche shift from the model by allowing adults and larvae to compete for and consume a shared resource, *K*. In the shared resource model the larval food equation and the adult fecundity terms are modified to account for the competition between the larvae and adults for the shared resource, with a constant *c* that determines the relative ability of larvae and adults to compete for food. The updated equations are:

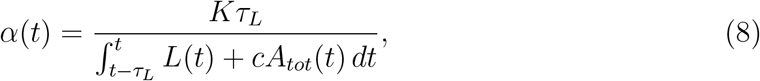

and

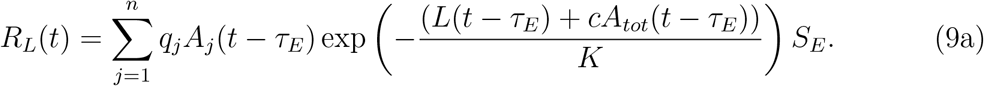

#### (3) Trait sensitivity model: Plasticity in only through-pupal survival

To isolate the effects of the capacity for trait plasticity, we limit plasticity to a single trait and in this simplified model we assume adult fecundity is not a plastic trait and does not vary with larval conditions; plasticity is only present in the trait, through-pupal survival, determined by a Holling type 3 function of larval food intake:

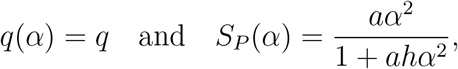

which only depends on two parameters: *a*, the consumption rate of the food and 1*/h*, the maximum value of through-pupal survival. By assuming fecundity is non-plastic we can isolate the effects of the degree of plasticity, sensitivity of a single trait to changes in the environment, on the existence of tipping points. Moreover, the Holling type 3 function gives a good fit to the through-pupal survival data, that is closely comparable to that obtained by Equation (6) (Figure 6), while also having the virtue of being described by only two parameters, allowing for a complete exploration of the effects of the degree of plasticity on population outcomes.

### 2.3 Defining the degree of plasticity (trait sensitivity)

For linear reaction norms, a common measure for the degree of plasticity, a measure of trait sensitivity to environmental change, is given by the reaction norm slope (Gómez *et al*., 2025), such that a steep slope corresponds to high trait plasticity and a shallow slope corresponds to low trait plasticity (Del Giudice, 2015). While nonlinear reaction norms are more difficult to characterise using a single metric, we examine two possible definitions of the degree of plasticity: i) the total number of trait responses an individual can express across all environments, and ii) the slope of the nonlinear reaction norm across a small range of environmental conditions (i.e., local plasticity). In the first of these definitions, when the number of trait responses (i.e., *n*, the number of size classes) is small these discrete traits are typically referred to as threshold traits, as the environmental cue must pass a threshold to elicit a trait change (Reid and Acker, 2022; Gómez *et al*., 2025). This is an example of polyphenism, whereby an individual takes on alternative discrete trait values in different environments, such as winged and wingless forms of aphids, or varying degrees of gregariousness in locus (Gómez *et al*., 2025). Except close to environmental thresholds, trait expression has little sensitivity to changes in the environment. At the other extreme, when *n* is large we have continuous trait change and a small change in environment is more likely to elicit a trait response. In the second definition of the degree of plasticity, we consider the trait sensitivity model which has a single continuous trait, through-pupal survival, that responds to changes in environment. The local slope of the reaction norm is increased by increasing either *a*, the consumption rate of the food, or 1*/h*, the maximum value of through-pupal survival. As with linear reaction norms, a steep local slope corresponds to a highly plastic trait, whereby a small change in environment results in a large trait change. We note that increasing *a* or 1*/h* has different effects on the overall reaction norm shape, despite both locally having similar effects on slope.

#### 2.3.1 Numerical simulations

In numerical simulations of the models, history for *t ≤* 0 is given by *L*(*t*) = 9500, *α*(*t*) = *K*_*L*_*/*9500, *A*_*i*_(*t*) = 0 for all *i* = 1, …, *n*. To simulate slowly decreasing ecosystem stress in Figure 3, *K*_*A*_ is initialised at 390 and simulations are run for 2000 days so that dynamics reach a stable state, *K*_*A*_ is then linearly increased from 390 to 450 at a rate of 0.001 per day, this is slow enough to allow dynamics to reach a stable state as the parameter is changed. For simulating slowly increasing ecosystem stress, *K*_*A*_ is decreased from *K*_*A*_ = 450 to *K*_*A*_ = 390 at the same rate of 0.001 per day. The model was simulated in Julia (Bezanson *et al*., 2017) using the package Differential Equations (Rackauckas and Nie, 2017). A table of model parameters is given in Supplementary Table S.1.1.

**Figure 3:**
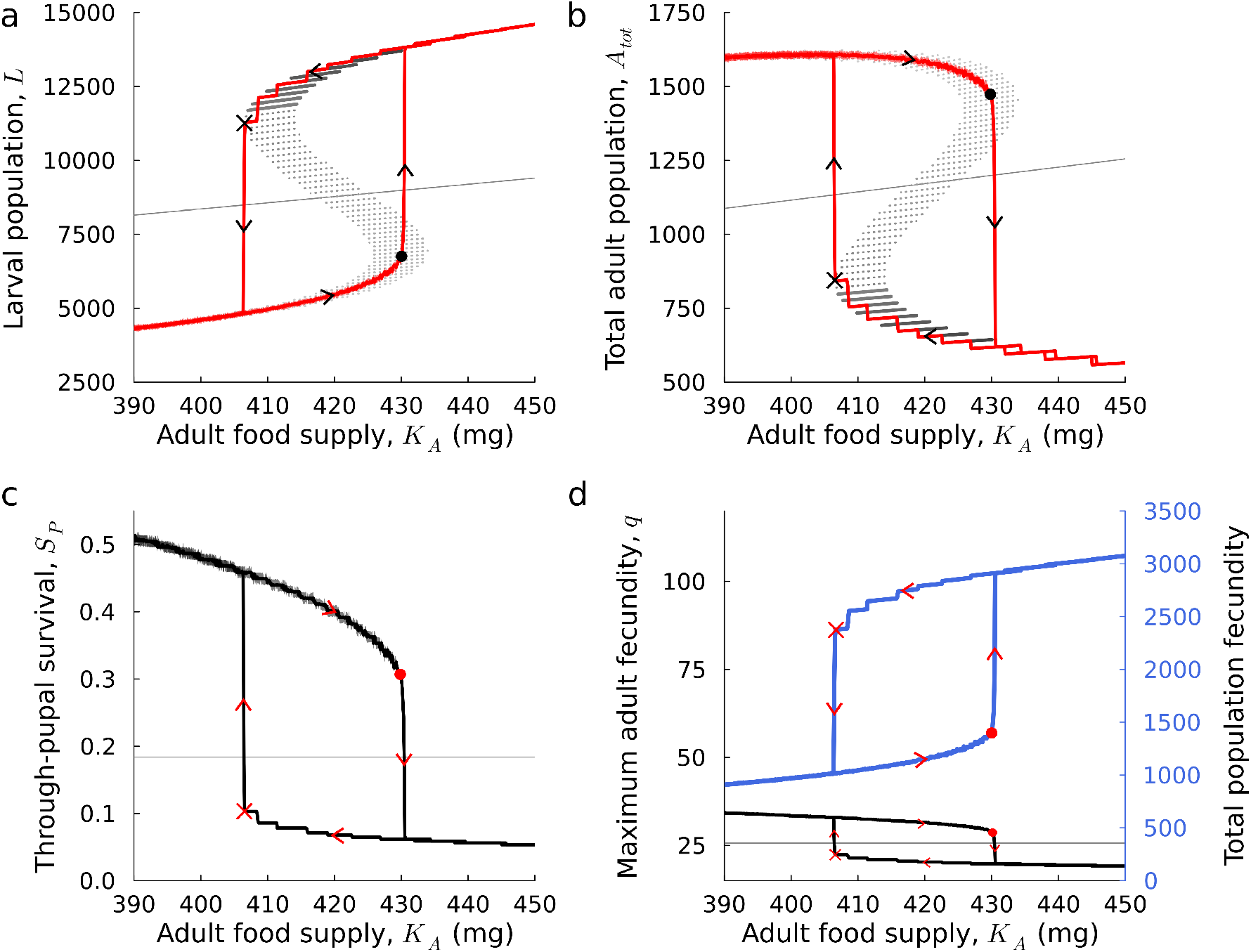
Varying adult food supply leads to tipping points induced by phenotypic plasticity, with critical transitions in abundance and traits. In the presence of phenotypic plasticity (main model), numerical solutions (red lines) for (**a**) the larval population and (**b**) the adult population demonstrate hysteresis as adult food *K*_*A*_, is slowly decreased and increased (indicated by the arrows). Tipping points occur at low (circle) and high (cross) levels of environmental stress. The black solid and dotted lines indicate, respectively stable and unstable analytical steady state solutions for each adult size class. The dotted lower branch in (**a**) and upper branch in (**b**) are associated with an unstable steady state, but this is accompanied by a stable periodic orbit around the steady state that maintains the branch. The traits (**c**) through-pupal survival and (**d**) maximum adult fecundity change as the daily supply of adult food is slowly increased and then decreased (indicated by the arrows). The blue line in (**d**) demonstrate how the total population fecundity (the number of eggs laid by the total population of adults in a day) undergoes critical transitions, in the opposing direction to the body-size related trait of maximum adult fecundity. In fixed trait model (simplified model (1)), *q* and *S*_*P*_ are constant (*q* = 25.66 and *S*_*P*_ = 0.18) and traits do not change as adult food *K*_*A*_ is varied, no hysteresis is observed, as indicated by the straight grey lines in (**a-d**). Model parameters are given in Table S.1.1, with *v*_3_ = *−*0.50.

To understand how plasticity influences the existence of tipping points, our analysis focusses on the effects of varying adult food supply, however our findings also hold when varying larval food supply instead. We focus on scenarios that give rise to population persistence and begin by using a steady state analysis (detailed in Supplementary S.2) in conjunction with numerical simulations to establish when alternative stable persistence steady states can occur and when bistability is lost. Loss of bistability results in a tipping point and a sudden change in population abundance. Using the various model simplifications we then isolate the potential drivers of this loss of bistability and determine the effects of varying the degree of plasticity on the existence and location of tipping points.

## 3 Results

### 3.1 Emergence of tipping points in response to changing adult food supply

The main blowfly model shows population persistence provided recruitment into the adult stage exceeds adult mortality, (*R*_0_(*α*) = *qS*_*E*_*S*_*L*_*S*_*P*_ (*α*)*S*_*J*_ */δ*_*A*_ *>* 1, the condition is derived in Supplementary S.2 and corresponds to the extinction steady state being unstable). At a low adult food supply (high extrinsic stress), the population settles to an adult dominated state comprised of a large adult population, with high values of fitness related traits (large adults with high survival and maximum fecundity) and a small larval population. Increasing the adult food supply, a low-stress tipping point (circles in Figure 3a-b) is crossed transitioning the system to one which is larval dominant, larval abundance is high, however both adult abundance and fitness related traits are low. Recovery to the original adult dominant state requires a significant reduction in the adult food supply, far in excess of the initial increase, to cross the high-stress tipping point (crosses in Figure 3a-b). The adult and larval dominated states represent alternative persistence steady states and tipping points occur via hysteresis when bistability is gained or lost, which is observed when slowly varying adult food supply, *K*_*A*_ (Figure 3a-b and full steady state analysis in Supplementary S.2). Sensitivity analysis (Supplementary S.3) demonstrates that tipping points are possible across a significant portion of parameter space.

The adult dominant state is maintained through small temporal oscillations in the population level trait distributions (Figure 4e), while the larval dominant state does not experience these oscillations and the population maintains a single shared phenotype. As larval abundance controls larval food per capita, critical transitions in traits occur along side critical transitions in population abundance (Figure 3c-d). The interaction between the abundance of each life-stage and the plastic life-history traits generates counterintuitive patterns in population abundance, namely that at high levels of adult food supply densities of adults can be low and the population dominated by larvae, in contrast at low levels of adult food supply the adults can dominate the population. Heuristically we can understand this pattern by noting that high adult food leads to low levels of adult competition, realised adult fecundity is therefore high (blue line in Figure 3d). The large birth rate results in high larval numbers and increased larval competition, leading to low trait values for through-pupal survival and maximum adult fecundity (constrains realised fecundity), that in turn lowers adult density in the next generation and instigates larval dominance. Analogous arguments apply to understand adult dominance. As the adult dominant state is also associated to larger adults than the larval dominant state (seen by the increased maximum adult fecundity in Figure 3d), the total biomass of the adult stage is significantly greater in the adult dominated state.

**Figure 4:**
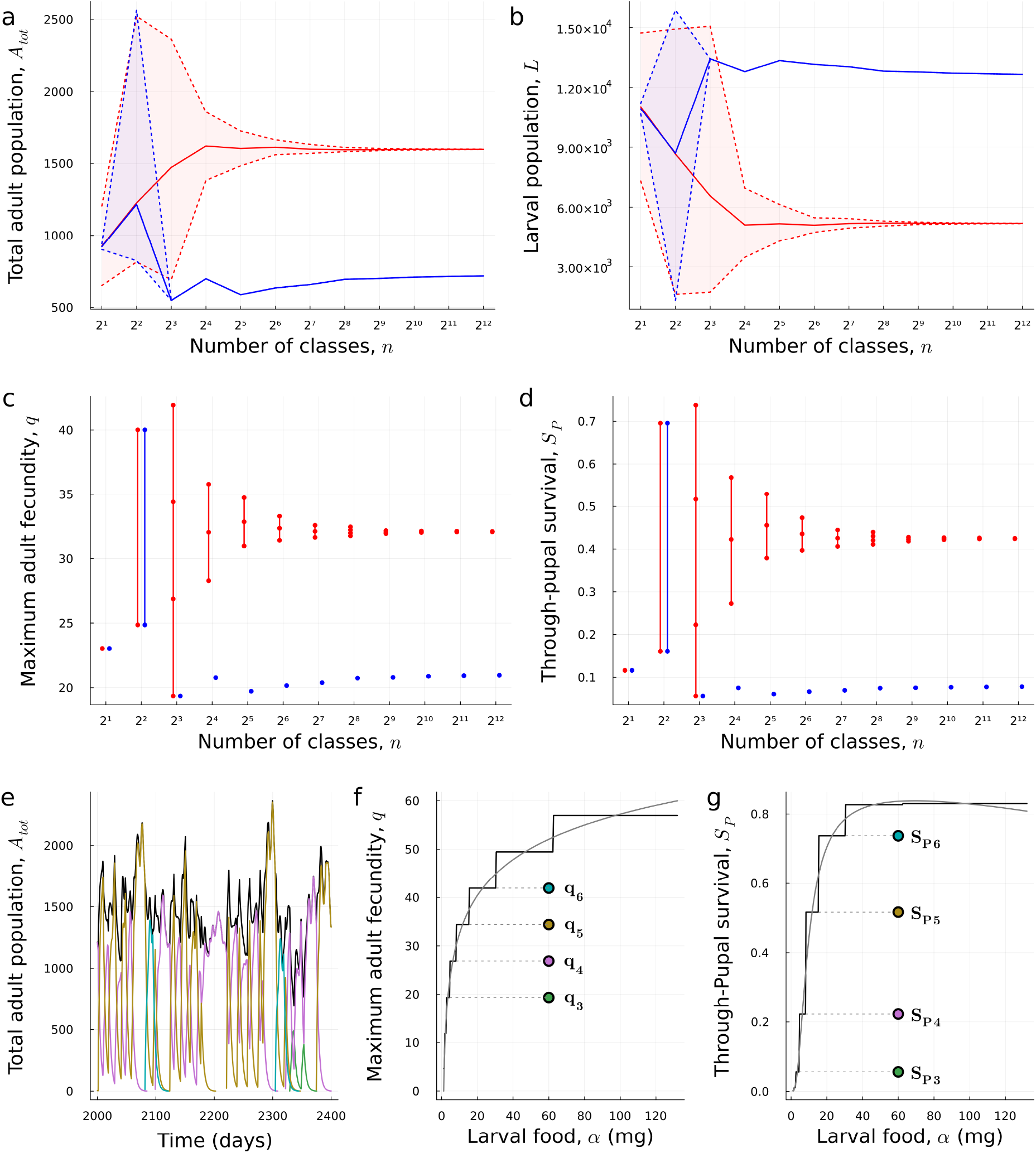
Varying the total number of trait responses to environmental change. Fixing *K*_*A*_ = 415 to ensure the existence of hysteresis, the number of adult trait classes (size classes), *n* is varied. The adult (**a**) and larval (**b**) abundance exhibit alternative stable states (red and blue lines and shading around the lines indicating the amplitude of population oscillations) when *n* is large (continuous traits), but hysteresis is lost when *n* very small (threshold traits). The population trait distribution shows a similar pattern as *n* is varied (**c**-**d**), where the dots indicate the traits found in the population. In (**e-g**) we illustrate the case of *n* = 2^3^, the number of trait classes is just sufficient to maintain the two alternative stable states. The stepped lines in (**f**) and (**g**) show the discretised reaction norms with 8 size classes and in (**e**) the temporal evolution of the adult population at the adult dominant state is plotted (black line), the coloured lines indicate the abundance of individuals with particular traits in the population, with the colours corresponding to the traits indicated in (**f**) and (**g**). The adult dominant state is maintained by a stable periodic orbit consisting of temporal fluctuations in the trait distribution of the population. The larval 1d3ominant state is not shown as no temporal changes in the trait distribution occur in this case and the single trait steady state is stable. Model parameters are given in Table S.1.1, with *K*_*A*_ = 415 and *v*_3_ = *−*0.50.

In consumer populations with complex lifecycles, like the blowfly system, the coupling of density-dependent feedbacks and ontogenetic niche shifts have been shown to induce tipping points between larval and adult dominated states (Schreiber and Rudolf, 2008). However, the fixed trait model (model simplification (1), when all individuals share a mean set of non-plastic traits), shows that hysteresis cannot occur in the absence of phenotypic plasticity, for any choice of trait values *q* and *S*_*P*_ (illustrated by the grey lines in Figure 3 and proved in general in Supplementary S.2.1). This demonstrates that an ontogenetic niche shift alone is not sufficient to induce tipping points and plasticity has a role to play. Similarly, in the shared resource model (model simplification (2), when adult and larvae consume a single shared resource, removing the ontogenetic niche shift) tipping points do not occur. Stability analysis of the shared resource model analytically proves that despite the presence of phenotypic plasticity, tipping points are not possible for any biologically reasonable choice of through-pupal survival and fecundity reaction norms (proved in general in Supplementary S.4). We can conclude that both an ontogenetic niche shift and phenotypic plasticity are needed to observe tipping points in the blowfly system.

The existence of tipping points in the main model is dependent on the shape of the reaction norms. A steady state analysis proves that convexity in the through-pupal survival reaction norm is a prerequisite to induce tipping points in the system (see analytical results in Supplementary S.2.1). For the blowfly, the convexity in the reaction norm is likely due to high competition for resources at high larval densities limiting size and consequently through-pupal survival, while at lower densities the competitive pressures are released and larvae can benefit from group feeding leading to a rapid increase in size and through-pupal survival (Moe *et al*., 2002). Convexity in through-pupal survival alone does not guarantee tipping points, rather the maximum adult fecundity reaction norm also plays a role, although a lesser one, in the induction of tipping points. Larger values and a lower gradient on the approach to the maximum adult fecundity aid in generating tipping points (Figure S.3.3b and Equation (S.2.4)). Given the importance of reaction norm shape in establishing phenotypic plasticity induced tipping points, in the next section we establish the robustness of this conclusion to variation in the degree of phenotypic plasticity, the sensitivity of trait responses to environmental change.

### 3.2 The degree of plasticity and strength of population feedbacks together determine the existence of tipping points

#### Reducing the total number of trait responses prevents tipping points

We found that reducing the degree of plasticity by decreasing the number of trait classes (*n*) (and therefore the total number of possible responses an individual can express in response to changes in resource availability), eliminated tipping points (Figure 4a-d). If traits are responsive enough to track very minor changes in environment (represented by lots of narrow trait classes, analogous to continuous traits), then we find tipping points. But when traits only respond to very large environmental changes (represented by a few wide trait classes, analogous to polyphenism), then tipping points are lost. This loss of tipping points occurs because the adult dominant state is maintained through small temporal oscillations in population level trait distribution; when traits have low sensitivity to changes in the environment (low *n*), the adult dominant branch cannot be maintained resulting in only a larval dominant state and no tipping points, regardless of the level of adult food supply.

By considering an example of 8 trait classes (Figure 4e-g), we see that the adult dominant state is maintained by trait switching, in this case only 4 out of the possible 8 traits are observed in the final adult dominant population. As the dominant population trait changes over time the population abundance responds, but as abundance influences trait expression via the reaction norms, trait sensitivity to changes in abundance is low when *n* is small and so population level traits do not immediately respond to the population change resulting in oscillations in abundance (light red shading in Figure 4a, as *n* decreases amplitude of oscillations increases). In the extreme case of very low trait sensitivity, the oscillations are of such large amplitude that they collide with the basin of attraction for the larval dominant state, at which point the adult dominant state can no longer be maintained (left end of Figure 4a and overlap of the light blue and red shaded areas).

#### Local plasticity alone does not determine tipping point existence

In this section we analyse the behaviour of the trait sensitivity model, which isolates the effects of local plasticity (in some small range of environmental conditions) in a single trait, through-pupal survival. The rationale for focussing on plasticity in a single trait comes from our finding that the shape of the through-pupal survival reaction norm determines the existence of tipping points. The degree of local plasticity in this trait is determined by varying the slope of the reaction norm, such that shallower reaction norm slopes correspond to smaller trait changes in response to environmental variation (less trait sensitivity). The local slope of the reaction norm can be reduced in one of two ways, by reducing the consumption rate (*a*) or reducing the maximum value of through-pupal survival (1*/h*) (Figure 5b and c). We prove that the trait sensitivity model continues to exhibit tipping points (analytical results in Supplementary S.5 and Figure 5a), which is illustrated by examining the larval and adult nullclines given by *dL/dt* = 0 and *dA/dt* = 0 (Figure 5d-e). Three intersections of the nullclines indicate the existence of alterative stable states. The adult nullcline (dotted line) is a decreasing function of *L* and the larval nullcline (solid line) is a unimodal function of *A* and so the nullclines typically intersect at one or three points, allowing for the possibility of alternative stable states. Both nullclines shift up and to the right as daily adult food supply (*K*_*A*_) is increased leading to the emergence of tipping points and hysteresis in this simplified system (Figure 5d). Changing daily larval food supply (*K*_*L*_) also leads to the emergence of tipping points. Changing larval food supply does not move the larval nullcline, but shifts the adult nullcline up and to the right creating alternative stables states in the process, these are then lost as larval food supply is increased further (Figure 5e and Supplementary S.5).

**Figure 5:**
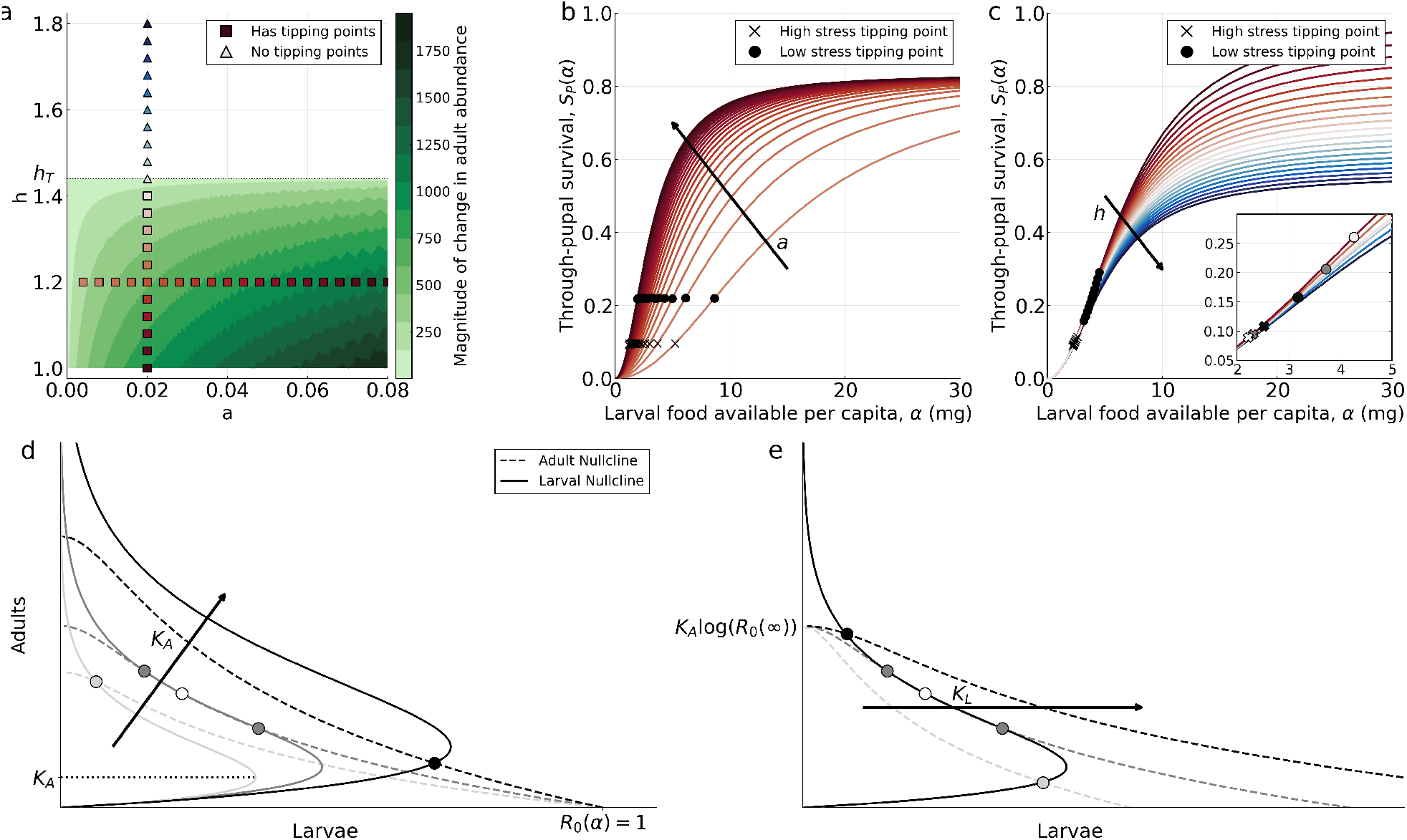
Effects of reaction norm shape on the existence of tipping points. The trait sensitivity model is considered, in which there is no plasticity in adult fecundity *q*(*α*) = 30 and through-pupal survival is a type 3 Holling function, *S*_*P*_ (*α*) = *aα*^2^*/*(1 + *ahα*^2^). (**a**), regions of *a*-*h* parameter space in which tipping points occur are indicated in green, with the shading reflecting the relative change in adult abundance between the two tipping points. Dark shading indicates large change in adult abundance as tipping points are crossed (strong hysteresis, large Δ*S*). Hysteresis occurs when *h* < *h*_*T*_ (see Supplementary S.5.3 for a derivation and expression for *h*_*T*_). The parameters *a* and 1*/h* control slope and maximum value of the reaction norm respectively. (**b)-(c**), are families of through-pupal survival reaction norms (*S*_*P*_ (*α*)) corresponding to the slices of two-dimensional parameter space indicated by the symbols in (**a**). In (**b**) *a* is varied, lighter red lines correspond to small values of *a* and darker red to high values. The location of the low and high stress tipping points are denoted by crosses and dots respectively. In (**c**) *h* is varied, lighter red lines correspond to small values of *h*, darker red lines correspond to large values of *h* and blue lines correspond very large values of *h* that fail to produce tipping points. In (**d**) and (**e**) the adult (dashed) and larval (solid) non-trivial nullclines are plotted showing how varying (**d**) *K*_*A*_ and (**e**) *K*_*L*_ result in the creation of alternative stable states. Filled circles indicate stable steady states and open circles indicate an unstable steady state. Light shading of the lines and filled circles correspond to low values of either *K*_*A*_ or *K*_*L*_, while darker shading correspond to higher values.

**Figure 6:**
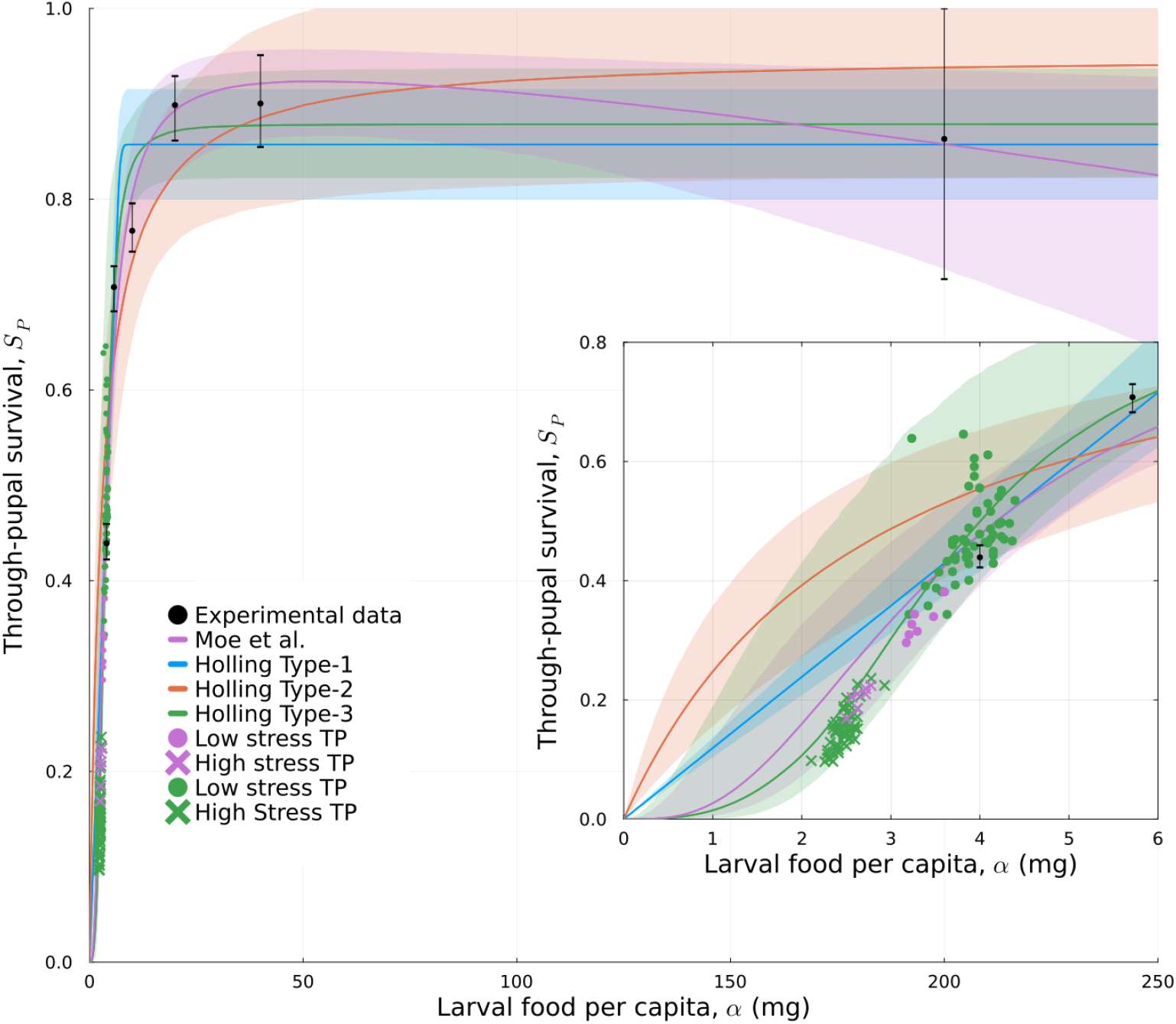
Comparison of four functional forms for the through-pupal survival reaction norm and their effect on tipping point induction. Four functional forms are fitted to the Moe *et al*. (2002) through-pupal survival data, these are the function of Moe *et al*. (2002) and three Holling functions (type 1, type 2, and type 3). Functions were fitted using the least mean square Julia package LsqFit, with residual sum of squares: 0.0804 (Moe *et al*.) and 0.0139, 0.0297, 0.00103 for types 1, 2 and 3, respectively and ΔAIC values of *−*0.72, 3.83, *−* 0.5214 respectively, when comparing to Moe *et al*.. For each functional form 4000 sample parameter sets were chosen from a multivariate normal distribution with the mean and covariance matrix obtained from fitting the function. The sample parameter sets give the median and 95% confidence intervals (shaded regions) for each functional form. For each functional form, 200 sample parameter sets were drawn from the 95% confidence intervals and the stage-structured model was simulated to identify if tipping points occurred, these were then indicated by the crosses (high environmental stress tipping point) and circles (low environmental stress tipping point).

In the trait sensitivity model plasticity only acts via through-pupal survival, consequently the strength of plasticity is entirely controlled by *a* (consumption rate) and 1*/h* (maximum value of through pupal survival). We prove that bistability only exists when the population level mean trait value of through-pupal survival is below 50% of the maximum through-pupal survival (*S*_*P*_ (*α*) < 1*/*2*h*, note that this condition is independent of *a*, see Supplementary S.5). Hysteresis additionally requires ln(*R*_0_(*α*)) *>* 1 (proved in Supplementary S.5), where *R*_0_(*α*) = *qS*_*E*_*S*_*L*_*S*_*P*_ (*α*)*S*_*J*_ */δ*_*A*_ is the expected number of progeny produced over an adult individual’s lifetime, this condition puts a lower bound on through-pupal survival. These two constraints mean that when tipping points exist they are located in regimes of low levels of larval food per capita in a region of the through-pupal survival reaction norm that exhibits a rapid increase in the survival proportion as larval food per capita is increased (Figure 5b-c). It has been suggested that tipping points in traits occur once species physiological limits have been reached, corresponding to reaction norm extrema (Cerini *et al*., 2023). Contrary to this theory, our evidence provides a counter-example to suggest tipping points can occur away from these extrema in traits and instead occur when trait expression is most sensitive to environmental change (Figure 5b-c).

Focussing in on the regime of low levels of larval food per capita, increasing *a* changes the slope of the reaction norm and hence the degree of plasticity and the sensitivity of traits to changes in environment. Increasing the steepness of the reaction norm (increasing *a*) does not change the population level mean trait value at the tipping point (no change in the vertical location of circles and crosses in Figure 5b). However, the steepness of the reaction norm does affect the magnitude of the response in population abundance at the tipping point (see definition of Δ*S* in Figure 1b and the darker green shading associated to high *a* in Figure 5a). A highly plastic trait (large *a* and locally steep reaction norm) is associated to a large population response and hence the strength of hysteresis is very strong for large *a*. The mechanism responsible for generating tipping points requires sufficient net population growth and hence through-pupal survival for density-dependent feedback to be strong enough to generate tipping points (ln(*R*_0_(*α*)) *>* 1), but if through-pupal survival is too high (*S*_*P*_ (*α*) *>* 1*/*2*h*) then there is only a very weak effect of larval density on population growth and the feedback mechanism is weakened. Hence, when *a* is large, to constrain survival within these bounds *α* and hence population density must be highly constrained and hysteresis, when it occurs, is necessarily very strong (see Supplementary S.5 for a proof of this result).

If instead we vary *h*, which determines the maximum value of the reaction norm, it does little to change the slope of the reaction norm in the range of *α* that gives rise to alternative stable states (see inset in Figure 5c). As *h* is increased the range of trait values that give rise to hysteresis reduces (see white to grey to back symbols in the inset in Figure 5c) until eventually hysteresis is no longer possible. Taking the local slope of the reaction norm as a measure of the degree of trait plasticity in this case leads to the conclusion that species with identical levels of trait plasticity can result in very different population dynamic outcomes, ranging from the existence of a tipping point to no tipping points.

## 4 Discussion

In a departure from current thinking (Stollewerk *et al*., 2025), we have shown how phenotypic plasticity, in response to resource availability, combined with population feedbacks can induce tipping points. This contrasts with the expectation that strong phenotypic plasticity reduces the risk of regime shifts by enabling a species to rapidly respond to environmental change (Stollewerk *et al*., 2025; Schneider, 2022). Instead, when an individual has a range of trait responses available to it, population level fluctuations in trait distribution enable the maintenance of alternative stable states allowing hysteresis to occur, while if trait plasticity is limited, the density-dependent feedbacks, coupled with an ontogenetic niche shift, that drive hysteresis are weakened, making regime shifts unlikely. In the extreme, in the absence of phenotypic plasticity, tipping points do not occur.

The mechanism driving the phenotypic plasticity induced tipping points is tightly linked to the complex life-cycle of the population, in particular the presence of an ontogenetic niche shift. The majority of species undergo a change in diet between life-stages (exhibit an ontogenetic niche shift) (Werner and Gilliam, 1984) and it has been demonstrated that this alone can generate hysteresis in response to changes in resource supply (Schreiber and Rudolf, 2008). The alternative stable states that are generated in this way are adult dominated or larvae dominated, as competition for resources in one life stage or the other acts as a bottleneck for recruitment to the next life-stage (Schröder *et al*., 2014). Plasticity in survival from larvae to adult regulates the bottleneck that determines adult abundance through the delayed response to changes in larval environment, while plasticity in fecundity regulates the bottleneck that determines juvenile abundance. The presence or absence of an ontogenetic niche shift substantially alters the relative strength of these bottlenecks. In this case, plasticity in survival is responsible for the existence of tipping points, but only when the relationship between survival and population density is convex. A linear survival reaction norm does not induce tipping points, even if the slope is large and there is a high degree of trait plasticity. A plausible mechanism for reaction norm convexity comes from larvae experiencing high levels of competition for food at high densities (low *α*), limiting size and survival, while at low densities competition is reduced and larvae can exploit group feeding allowing a steep increase in survival (Moe *et al*., 2002). This rapid change in survival provides a source of strong density-dependent feedback that can main alternative state states.

Reaction norms are routinely used to infer the effects of environmental change on population fitness (as discussed by Brass *et al*. (2021)), but even when the traits are fitness related, increases in individual fitness traits do not always carry through to increases in population fitness (Mclean *et al*., 2016; Iler *et al*., 2021). We find an extreme example of this when survival reaction norms with almost identical slopes locally over a range of environmental conditions, give rise to very different population responses to environmental change, ranging from critical regimes shifts to almost no population change. These different outcomes are due to the maximum range of possible trait responses defined by the reaction norm in each case differing. Alternative stable states are maintained when survival distributions are below half of the maximum survival, higher than this there is no strong bottle neck to generate the alternative stable states. However, if the maximum survival is low, any rapid increase in survival that comes with release from competition, is too small to generate the strong feedback required to maintain alternative stable states. So differences in outcome are related to the full reaction norm and not just responses to a narrow range of environmental conditions. This can have implications for data col-lection, as we see in the blowfly example, as the tipping points were found to occur in regimes of low through-pupal survival, where there can be a lack of experimental data (< 4mg of larval food per capita in Figure 6) to distinguish reaction norm shape. By comparing Moe *et al*. (2002) through-pupal survival reaction norm, with three plausible alternatives, namely the Holling type 1, 2 and 3 functions (Holling, 1959), that all fit the data well, only the Moe *et al*. (2002) and the Holling type 3 functions satisfy the convexity condition and lead to the existence of tipping points in the population model (Figure 6). These findings highlight the importance of considering the full range of environmental conditions when constructing reaction norms to predict population outcomes, as trait plasticity is propagated and amplified through the trait-population feedbacks leading to unexpected consequences at the population level.

It has been hypothesised that early warning signals of regime-shift may exist as a series of predictable sequential shifts in species traits, in response to environmental change prior to a demographic response (Cerini *et al*., 2023; Clements and Ozgul, 2018; Baruah *et al*., 2020). The predictability comes from traits reaching their physiological limits. However, we have found that the population trait distributions at tipping points can be concentrated well away from the limit of the individual-level plastic capacity. The reaction norms point to further trait adaptation being possible under increased environmental stress, but interactions with population dynamics prevent the additional trait adaptation occurring, due to abundance constraining the traits that are can be expressed. This poses additional challenges for the search for reliable early warning signals for catastrophic ecosystem regime shifts.

While our work has been to consider a single environmental stressor and a simple set of traits, species often have many plastic life-history traits and phenotypic plasticity can be a response to multiple environmental stressors (Forsman, 2015; Couret *et al*., 2014; Westneat *et al*., 2019). A more complex network of traits and environmental stressors provides additional opportunities for the types of positive feedback that commonly cause hysteresis and tipping points. However, these additional feedbacks could also act to temper the mechanisms driving regime shifts that have been discussed here (Mclean *et al*., 2016). Therefore, a whole population and organism approach is needed to effectively unpick the relationship between tipping points and phenotypic plasticity. One such approach is to use dynamic energy budget theory to track the flow of energy allocation from the individual all the way up to the population level response (Croll and de Roos, 2022; Smallegange *et al*., 2017).

Beyond considering individual species a recent review highlights that the feedbacks we discuss here also operate across higher levels of organisation, to communities (Stollewerk *et al*., 2025). Our theoretical approach could readily be extended to interacting species and provides an adaptable framework for unpicking the effects of more complex networks of traits and stressors than those studied here. Moreover, it is amenable to analysis, offering a potential route to mechanistically understand the role of phenotypic plasticity in regime shifts in a systematic way. Understanding the complex interplay between phenotypic plasticity, environmental stress, and population dynamics is essential for predicting and mitigating biodiversity loss. Our findings highlight that while plasticity can sometimes buffer against environmental change, it can also unexpectedly drive populations toward tipping points, resulting in devastating and catastrophic ecosystem collapse. Species with traits that respond to changes in resource availability or crowding may be particularly vulnerable to plasticity induced collapse as these traits provide a route for the density-dependent feedbacks that can generate hysteresis at the population level (Kéfi *et al*., 2016). However, refining our models and incorporating biologically informed reaction norms, presents an opportunity to improve our ability to anticipate these shifts, and a whole-population approach that considers multiple stressors and complex life histories will be critical in safeguarding ecosystems through targeted conservation strategies. By working towards more accurate predictions and effective solutions, we provide hope that science can guide interventions to preserve biodiversity in an era of rapid environmental change.

## Acknowledgements

The authors would like to thank two anonymous referees for many helpful suggestions that improved the quality of this article. BF was funded by an EPSRC DTP studentship 2608237. CC, SW, DB and AN were supported by EPSRC grants EP/Y017838/1 and EP/Y017919/1.

## Author contributions

CC, SW and BF conceptualised the study and methodology. BF, developed the methodology, conducted the analysis developed the code, under the supervision of CC, SW and DB. BF, CC wrote the paper, SW, DB, AN reviewed and edited the manuscript.

## Data Availability Statement

The data that support the findings of this study are openly available on Zenodo at http://doi.org/10.5281/zenodo.15675297, reference number 15675297.

## Conflict of Interest Statement

The authors declare no conflicts of interest.

## Notes

http://doi.org/10.5281/zenodo.15675297

### Competing Interest Statement

The authors have declared no competing interest.

