## Supplementary information for "Phenotypic plasticity as a route to population shifts via tipping points"

### S.1 Model parameter values

| Parameter | Biological interpretation | Value | Parameter source |
|---|---|---|---|
| $\tau_E$ | Egg stage duration (days) | 0.6 | Gurney *et al.* (1983) |
| $\tau_L$ | Larval stage duration (days) | 5 | Gurney *et al.* (1983) |
| $\tau_P$ | Pupal stage duration (days) | 5.9 | Gurney *et al.* (1983) |
| $\tau_J$ | Juvenile adult stage duration (days) | 4.1 | Gurney *et al.* (1983) |
| $\delta_E$ | Per capita egg mortality rate (day$^{-1}$) | 0.07 | Gurney *et al.* (1983) |
| $\delta_L$ | Per capita larval mortality rate (days$^{-1}$) | 0.004 | Gurney *et al.* (1983) |
| $\delta_J$ | Per capita juvenile adult mortality rate (days$^{-1}$) | 0.0025 | Gurney *et al.* (1983) |
| $\delta_A$ | Per capita reproductive-adult mortality rate (days$^{-1}$) | 0.27 | Gurney *et al.* (1983) |
| $K_L$ | Daily larval food supply in the Nicholson experiments (mg of protein) | 50000 | Nicholson (1957) |
| $K_A$ | Daily supply of adult food (mg) | Varied | N/A |
| $v_1$ | Fitted constant in the through-pupal survival reaction norm | $-4.00 \pm 1.49$ | Moe *et al.* (2002) |
| $v_2$ | Fitted constant in the through-pupal survival reaction norm | $3.36 \pm 0.59$ | Moe *et al.* (2002) |
| $v_3$ | Fitted constant in the through-pupal survival reaction norm | $-0.44 \pm 0.06$ | Moe *et al.* (2002) |
| $K_{LM}$ | Daily larval food supply in the Moe et al. experiments (mg of protein) | 2000 | Brass *et al.* (2021); Moe *et al.* (2002) |
| $h_1$ | Fitted constant in the adult fecundity reaction norm | 3.95 | Webber (1955) |
| $h_2$ | Fitted constant in adult fecundity reaction norm | 6.90 | Webber (1955) |
| $h_3$ | Fitted constant in the adult fecundity reaction norm | 0.97 | Webber (1955) |
| $h_4$ | Fitted constant in the adult fecundity reaction norm | 0.78 | Webber (1955) |
| $q_{max}$ | Highest maximum adult fecundity | 60 | Webber (1955) |
| $q_{min}$ | Lowest maximum adult fecundity | 0 | Selected |
| $n$ | Number of adult size classes | 256 | Selected |

Table S.1.1: **Model parameter values** For parameter values characterising the through-pupal survival taken from Moe *et al.* (2002), one standard error from the mean values is also given, standard deviations were not available for other model parameter estimates. In contrast to Moe *et al.*, who considered limited larval food (2000mg of protein per day), the Nicholson experiments provided the larvae with unlimited food (50000 mg of protein per day) and we consider the unlimited food scenario in the model simulations and analysis.

### S.2 Model stability analysis

The dynamical behaviour of the Brass *et al.* (2021) model (Equations 1) can be classified by a stability analysis. There are two classes of steady state, (i) the extinction state given by $L(t) = 0$, $A_i(t) = 0$ for all $i$, with $\alpha(t)$ a constant determined by initial conditions, and (ii) a family of persistence steady states given by $\alpha(t) = \alpha^*$, $L(t) = L^*$, $A_k(t) = A_k^*$ and $A_i(t) = 0$ for all $i \neq k$, where all adults occupy a single size-class. The size class $k$ and the plastic traits of individuals are determined by the value of $\alpha^* \in (\alpha_{k-1}, \alpha_k)$. Adults occupy a single size class because $\omega_i(\alpha^*)$ is either 0 or 1, so all maturing larvae transition into a single adult size class.

Solving Equations 1 to find the persistence steady state equations gives:

$$\alpha^* = \frac{K_L \delta_L S_L S_{Pk} S_J}{K_A \delta_A (1 - S_L) \ln\left(\frac{q_k S_E S_L S_{Pk} S_J}{\delta_A}\right)}, \tag{S.2.1a}$$

$$L^* = \frac{K_A \delta_A (1 - S_L)}{\delta_L S_L S_{Pk} S_J} \ln\left(\frac{q_k S_E S_L S_{Pk} S_J}{\delta_A}\right), \tag{S.2.1b}$$

$$A_k^* = K_A \ln\left(\frac{q_k S_E S_L S_{Pk} S_J}{\delta_A}\right), \tag{S.2.1c}$$

$$A_i^* = 0 \text{ for } i = 1, ..., k-1, k+1, ..., n. \tag{S.2.1d}$$

The persistence steady state is biologically feasible (non-negative) provided $q_k S_E S_L S_{Pk} S_J > \delta_A$, which states that the recruitment rate into adult size-class $k$ is larger than the adult mortality rate. In Section S.2.2 we show that this condition on adult recruitment and survival is precisely the condition that ensures the extinction steady state is unstable and the population persists.

#### S.2.1 Existence of alternative persistence steady states

Equations S.2.1 can have multiple solutions, giving rise to more than one persistent steady state in which adults occupy the single adult size-class $k$. To establish the existence of multiple solutions it is useful to express the steady state equations in terms of $S_P(\alpha^*)$ and $q(\alpha^*)$ rather than the discretised terms $S_{Pk}$ and $q_k$. We can then solve Equations S.2.1 for the bifurcation parameter, $K_A$, the daily adult food supplied:

$$K_A = \frac{K_L \delta_L S_L S_P(\alpha^*) S_J}{\alpha^* \delta_A (1 - S_L) \ln\left(\frac{q(\alpha^*) S_E S_L S_P(\alpha^*) S_J}{\delta_A}\right)}. \tag{S.2.2}$$

The existence of multiple solutions, $\alpha^*$, to Equation (S.2.2) is a requirement for the existence of tipping points. Multiple solutions exist if Equation (S.2.2) has at least one turning point. Differentiating $K_A$ with respect to $\alpha^*$ gives:

$$\frac{dK_A}{d\alpha^*} = K_A \left( \frac{S_P'(\alpha^*)}{S_P(\alpha^*)} - \frac{1}{\alpha^*} - \frac{1}{\ln\left(\frac{q(\alpha^*) S_E S_L S_P(\alpha^*) S_J}{\delta_A}\right)} \left( \frac{q'(\alpha^*)}{q(\alpha^*)} + \frac{S_P'(\alpha^*)}{S_P(\alpha^*)} \right) \right). \tag{S.2.3}$$

Given $K_A > 0$, turning points of Equation (S.2.2) satisfy:

$$\frac{S_P'(\alpha^*)}{S_P(\alpha^*)} - \frac{1}{\alpha^*} = \frac{1}{\ln\left(\frac{q(\alpha^*) S_E S_L S_P(\alpha^*) S_J}{\delta_A}\right)} \left( \frac{q'(\alpha^*)}{q(\alpha^*)} + \frac{S_P'(\alpha^*)}{S_P(\alpha^*)} \right). \tag{S.2.4}$$

To study the solutions of Equation (S.2.4), we define:

$$f(\alpha) = \frac{S_P'(\alpha)}{S_P(\alpha)} - \frac{1}{\alpha}, \tag{S.2.5a}$$

$$g(\alpha) = \frac{1}{\ln\left(\frac{q(\alpha) S_E S_L S_P(\alpha) S_J}{\delta_A}\right)} \left( \frac{q'(\alpha)}{q(\alpha)} + \frac{S_P'(\alpha)}{S_P(\alpha)} \right). \tag{S.2.5b}$$

We first note that if there is no plasticity in either trait (i.e. through-pupal survival and maximum adult fecundity do not change with the average daily per capita larval food), then $g(\alpha) = 0$ and $f(\alpha) < 0$ yielding $dK_A/d\alpha < 0$ for all $\alpha$ and there is only one persistence steady state corresponding to each adult size class and tipping points are not possible.

Provided the persistence steady state exists ($q S_E S_L S_P(\alpha) S_J > \delta_A$), then when through-pupal survival and maximum adult fecundity increase as a function of per capita larval food $g(\alpha) > 0$, we have $f(\alpha) > 0$ is a necessary condition for Equation (S.2.2) to have multiple solutions. We can show that the condition $f(\alpha) > 0$ requires the through-pupal survival function to be convex in some range of $\alpha$.

Suppose that there exists $\alpha^* = \alpha_a$ such that $f(\alpha_a) > 0$, then $S'_P(\alpha_a) > S_P(\alpha_a)/\alpha_a$. By the mean value theorem (MVT) there exists an $\alpha_b \in (0, \alpha_a)$ such that

$$S'_P(\alpha_b) = \frac{S_P(\alpha_a) - S_P(0)}{\alpha_a - 0} \leq \frac{S_P(\alpha_a)}{\alpha_a} < S'_P(\alpha_a), \tag{S.2.6}$$

where the first inequality is due to $S_P(0) \geq 0$ and the second inequality is by the assumption. This gives a condition on the first derivative of $S_P$, however, in order to show convexity, we need the second derivative of $S_P(\alpha)$ to be positive. Applying the MVT to $S'_P(\alpha)$, there exists an $\alpha_c \in (\alpha_b, \alpha_a)$ such that

$$S''_P(\alpha_c) = \frac{S'_P(\alpha_a) - S'_P(\alpha_b)}{\alpha_a - \alpha_b} > 0, \tag{S.2.7}$$

where the inequality is due to Equation (S.2.6). Hence $S_P(\alpha)$ is convex in some region around $\alpha_c$. Convexity alone is not enough to guarantee the existence of hysteresis and tipping points, but is a requirement. In order for hysteresis to occur, there must be at least two turning points and $f(\alpha) = g(\alpha)$ requires two solutions. Satisfying this additional requirement depends on the properties of both reaction norms, $S_P(\alpha)$ and $q(\alpha)$.

So far we have assumed through-pupal survival is a strictly increasing function of the average daily amount of larval food per capita, while this is a reasonable assumption, it is worth considering the other cases. If we consider the case in which through-pupal survival is not a plastic trait ($S'_P(\alpha) = 0$), then Equation (S.2.4) becomes

$$q'(\alpha^*) = -\frac{q(\alpha^*)}{\alpha^*} \ln\left(\frac{q(\alpha^*) S_E S_L S_P S_J}{\delta_A}\right) \tag{S.2.8}$$

Since $q(\alpha^*) S_E S_L S_P S_J > \delta_A$ is required for population persistence, Equation (S.2.8) implies that the requirement for the existence of tipping points when there is no plasticity in through-pupal survival, is $q'(\alpha^*) < 0$ and maximum fecundity must decline with average larval food per capita for some range of $\alpha$.

Finally, we consider the case where $q'(\alpha) > 0$, but $S'_P(\alpha) < 0$ in some range of $\alpha$. This can occur if there is an Allee effect, for example, if early instar larvae benefit from more efficient feeding in large groups, resulting in higher through-pupal survival at low $\alpha$ (Moe *et al.*, 2002). In this scenario, we rearrange Equation (S.2.4) to arrive at $F(\alpha) = G(\alpha)$, where

$$F(\alpha) = \frac{S'_P(\alpha)}{S_P(\alpha)} \left(1 - \frac{1}{\ln\left(\frac{q(\alpha) S_E S_L S_P(\alpha) S_J}{\delta_A}\right)}\right), \tag{S.2.9a}$$

$$G(\alpha) = \frac{1}{\ln\left(\frac{q(\alpha) S_E S_L S_P(\alpha) S_J}{\delta_A}\right)} \frac{q'(\alpha)}{q(\alpha)} + \frac{1}{\alpha}. \tag{S.2.9b}$$

As $G(\alpha) > 0$, a necessary condition for tipping points is $F(\alpha) > 0$, or equivalently

$$\delta_A < q(\alpha) S_E S_L S_P(\alpha) S_J < \exp(1)\delta_A. \tag{S.2.10}$$

The above constraint restricts the parameters to a very small region of parameter space, making the existence of tipping points unlikely when $S'_P(\alpha) < 0$.

#### S.2.2 Stability of the extinction steady state

In this section, we demonstrate that the extinction steady state is stable provided $q(\alpha^*) S_E S_L S_P(\alpha^*) S_J < \delta_A$. Without loss of generality, we assume $\omega_1(\alpha^*) = 1$ and $\omega_i(\alpha^*) = 0$ for $i \neq 1$. We introduce $\bar{\alpha}(t) = \alpha^* + \alpha(t)$, $\bar{L}(t) = 0 + L(t)$, and $\bar{A}_i(t) = 0 + A_i(t)$ and linearise Equations 1 around the extinction steady steady state to give:

$$
\begin{aligned}
\begin{pmatrix} \frac{d\bar{\alpha}}{dt}(t) \\ \frac{d\bar{L}}{dt}(t) \\ \frac{d\bar{A}_1}{dt}(t) \\ \frac{d\bar{A}_2}{dt}(t) \\ \vdots \\ \frac{d\bar{A}_n}{dt}(t) \end{pmatrix} = V_0 \begin{pmatrix} \bar{\alpha}(t) \\ \bar{L}(t) \\ \bar{A}_1(t) \\ \bar{A}_2(t) \\ \vdots \\ \bar{A}_n(t) \end{pmatrix} + V_L \begin{pmatrix} \bar{\alpha}(t-\tau_L) \\ \bar{L}(t-\tau_L) \\ \bar{A}_1(t-\tau_L) \\ \bar{A}_2(t-\tau_L) \\ \vdots \\ \bar{A}_n(t-\tau_L) \end{pmatrix} + V_E \begin{pmatrix} \bar{\alpha}(t-\tau_E) \\ \bar{L}(t-\tau_E) \\ \bar{A}_1(t-\tau_E) \\ \bar{A}_2(t-\tau_E) \\ \vdots \\ \bar{A}_n(t-\tau_E) \end{pmatrix} \\
+ V_{EL} \begin{pmatrix} \bar{\alpha}(t-\tau_E-\tau_L) \\ \bar{L}(t-\tau_E-\tau_L) \\ \bar{A}_1(t-\tau_E-\tau_L) \\ \bar{A}_2(t-\tau_E-\tau_L) \\ \vdots \\ \bar{A}_n(t-\tau_E-\tau_L) \end{pmatrix} + V_{ELPJ} \begin{pmatrix} \bar{\alpha}(t-\tau) \\ \bar{L}(t-\tau) \\ \bar{A}_1(t-\tau) \\ \bar{A}_2(t-\tau) \\ \vdots \\ \bar{A}_n(t-\tau) \end{pmatrix},
\end{aligned}
\tag{S.2.11}
$$

where $\tau = \tau_E + \tau_L + \tau_P + \tau_J$, and

$$
V_0 = \begin{pmatrix} 0 & -(\alpha^*)^2/(K_L\tau_L) & 0 & 0 & \cdots & 0 \\ 0 & -\delta_L & 0 & 0 & \cdots & 0 \\ 0 & 0 & -\delta_A & 0 & \cdots & 0 \\ 0 & 0 & 0 & -\delta_A & \cdots & 0 \\ \vdots & \vdots & \vdots & \vdots & \ddots & \vdots \\ 0 & 0 & 0 & 0 & \cdots & -\delta_A \end{pmatrix}, V_L = \begin{pmatrix} 0 & (\alpha^*)^2/(K_L\tau_L) & 0 & 0 & \cdots & 0 \\ 0 & 0 & 0 & 0 & \cdots & 0 \\ 0 & 0 & 0 & 0 & \cdots & 0 \\ 0 & 0 & 0 & 0 & \cdots & 0 \\ \vdots & \vdots & \vdots & \vdots & \ddots & \vdots \\ 0 & 0 & 0 & 0 & \cdots & 0 \end{pmatrix},
$$

$$
V_E = \begin{pmatrix} 0 & 0 & 0 & 0 & \cdots & 0 \\ 0 & 0 & q_1S_E & q_2S_E & \cdots & q_nS_E \\ 0 & 0 & 0 & 0 & \cdots & 0 \\ 0 & 0 & 0 & 0 & \cdots & 0 \\ \vdots & \vdots & \vdots & \vdots & \ddots & \vdots \\ 0 & 0 & 0 & 0 & \cdots & 0 \end{pmatrix}, V_{EL} = \begin{pmatrix} 0 & 0 & 0 & 0 & \cdots & 0 \\ 0 & 0 & -q_1S_ES_L & -q_2S_ES_L & \cdots & -q_nS_ES_L \\ 0 & 0 & 0 & 0 & \cdots & 0 \\ 0 & 0 & 0 & 0 & \cdots & 0 \\ \vdots & \vdots & \vdots & \vdots & \ddots & \vdots \\ 0 & 0 & 0 & 0 & \cdots & 0 \end{pmatrix},
$$

$$
V_{ELPJ} = \begin{pmatrix} 0 & 0 & 0 & 0 & \cdots & 0 \\ 0 & 0 & 0 & 0 & \cdots & 0 \\ 0 & 0 & q_1S_ES_LS_{P1}S_J & q_2S_ES_LS_{P1}S_J & \cdots & q_nS_ES_LS_{P1}S_J \\ 0 & 0 & 0 & 0 & \cdots & 0 \\ \vdots & \vdots & \vdots & \vdots & \ddots & \vdots \\ 0 & 0 & 0 & 0 & \cdots & 0 \end{pmatrix}.
$$

The matrices $V_0, V_E, V_L, V_{EL}, V_{ELPJ}$ are upper triangular and hence the characteristic equation for Equation (S.2.11) simplifies to:

$$
\lambda(\lambda+\delta_L)(\lambda+\delta_A)^{n-1}\left(\lambda+\delta_A - q_1S_ES_LS_{P1}S_Je^{-\tau\lambda}\right) = 0. \tag{S.2.13}
$$

The eigenvalues are $\lambda = 0$, $\lambda = -\delta_L$, $\lambda = -\delta_A$ and the solution to

$$
\lambda = -\delta_A + q_1S_ES_LS_{P1}S_Je^{-\tau\lambda}. \tag{S.2.14}
$$

The last equation has a countably infinite number of solutions (which includes complex solutions), however, if the solution with largest real part has positive real part the extinction steady state is unstable.

Theorem 4.7 of Smith (2011) states that given $\lambda = A + Be^{-\lambda r}$, where $A, B, r \in \mathbb{R}$, the solution, $\lambda$, with largest real part satisfies (a) $Re(\lambda) > 0$ if $A + B > 0$ and (b) $Re(\lambda) < 0$ if $A + B < 0$ and $B \geq A$. Applying this theorem to Equation (S.2.14) we conclude that the extinction steady state is unstable when $q(\alpha^*)S_ES_LS_P(\alpha^*)S_J > \delta_A$, that is, when the recruitment rate into the adult stage is greater than the adult mortality rate. When $q(\alpha^*)S_ES_LS_P(\alpha^*)S_J < \delta_A$, the extinction state is the only biologically feasible equilibrium, and hence despite the presence of the zero eigenvalue (due to the neutral stability to perturbations in $\alpha$), we expect the extinction steady state to be stable. Defining the lifetime reproductive potential as $R_0(\alpha^*) = q(\alpha^*)S_ES_LS_P(\alpha^*)S_J/\delta_A$, we have population persistence when $R_0(\alpha^*) > 1$.

#### S.2.3 Stability of the persistence steady state(s)

To analyse the stability of the persistence steady state we first note that without a loss of generality we can assume all adults occupy size class $i = 1$, such that $\omega_1(\alpha^*) = 1$ and $\omega_i(\alpha^*) = 0$ for $i \neq 1$ and the first adult size-class is non-zero at steady state and $\alpha^* \in (\alpha_0, \alpha_1)$. Since each function $\omega_i(\alpha)$ takes the value of either 1 or 0, for $\alpha$ within the interior of a size class there is a small region around $\alpha$ such that $\partial\omega_i/\partial\alpha = 0$. This implies provided $\alpha^*$ is away from a class boundary then $\partial\omega_i/\partial\alpha(\alpha^*) = 0$ for each $i$, assuming this is the case, we introduce $\bar{\alpha}(t) = \alpha^* + \alpha(t)$, $\bar{L}(t) = L^* + L(t)$, $\bar{A}_1(t) = A_1^* + A_1(t)$, and $\bar{A}_i(t) = 0 + A_i(t)$ for $i = 2, 3, \ldots, n$ and linearise Equations 1 around the persistence steady state, in which all adults are in size class 1, yielding:

$$
\begin{aligned}
\begin{pmatrix} \frac{d\bar{\alpha}}{dt}(t) \\ \frac{d\bar{L}}{dt}(t) \\ \frac{d\bar{A}_1}{dt}(t) \\ \frac{d\bar{A}_2}{dt}(t) \\ \vdots \\ \frac{d\bar{A}_n}{dt}(t) \end{pmatrix} &= V_0 \begin{pmatrix} \bar{\alpha}(t) \\ \bar{L}(t) \\ \bar{A}_1(t) \\ \bar{A}_2(t) \\ \vdots \\ \bar{A}_n(t) \end{pmatrix} + V_L \begin{pmatrix} \bar{\alpha}(t-\tau_L) \\ \bar{L}(t-\tau_L) \\ \bar{A}_1(t-\tau_L) \\ \bar{A}_2(t-\tau_L) \\ \vdots \\ \bar{A}_n(t-\tau_L) \end{pmatrix} + V_E \begin{pmatrix} \bar{\alpha}(t-\tau_E) \\ \bar{L}(t-\tau_E) \\ \bar{A}_1(t-\tau_E) \\ \bar{A}_2(t-\tau_E) \\ \vdots \\ \bar{A}_n(t-\tau_E) \end{pmatrix} \\
&\quad + V_{EL} \begin{pmatrix} \bar{\alpha}(t-\tau_E-\tau_L) \\ \bar{L}(t-\tau_E-\tau_L) \\ \bar{A}_1(t-\tau_E-\tau_L) \\ \bar{A}_2(t-\tau_E-\tau_L) \\ \vdots \\ \bar{A}_n(t-\tau_E-\tau_L) \end{pmatrix} + V_{ELPJ} \begin{pmatrix} \bar{\alpha}(t-\tau) \\ \bar{L}(t-\tau) \\ \bar{A}_1(t-\tau) \\ \bar{A}_2(t-\tau) \\ \vdots \\ \bar{A}_n(t-\tau) \end{pmatrix},
\end{aligned}
\tag{S.2.15}
$$

where $\tau = \tau_E + \tau_L + \tau_P + \tau_J$ and

$$
V_0 = \begin{pmatrix} 0 & -(\alpha^*)^2/(K_L\tau_L) & 0 & 0 & \cdots & 0 \\ 0 & -\delta_L & 0 & 0 & \cdots & 0 \\ 0 & 0 & -\delta_A & 0 & \cdots & 0 \\ 0 & 0 & 0 & -\delta_A & \cdots & 0 \\ \vdots & \vdots & \vdots & \vdots & \ddots & \vdots \\ 0 & 0 & 0 & 0 & \cdots & -\delta_A \end{pmatrix}, V_L = \begin{pmatrix} 0 & (\alpha^*)^2/(K_L\tau_L) & 0 & 0 & \cdots & 0 \\ 0 & 0 & 0 & 0 & \cdots & 0 \\ 0 & 0 & 0 & 0 & \cdots & 0 \\ 0 & 0 & 0 & 0 & \cdots & 0 \\ \vdots & \vdots & \vdots & \vdots & \ddots & \vdots \\ 0 & 0 & 0 & 0 & \cdots & 0 \end{pmatrix},
$$

$$
V_E = \begin{pmatrix} 0 & 0 & 0 & 0 & \cdots & 0 \\ 0 & 0 & q_1(1 - A_1^*/K_A)e^{-A_1^*/K_A}S_E & (q_2 - q_1A_1^*/K_A)e^{-A_1^*/K_A}S_E & \cdots & (q_n - q_1A_1^*/K_A)e^{-A_1^*/K_A}S_E \\ 0 & 0 & 0 & 0 & \cdots & 0 \\ 0 & 0 & 0 & 0 & \cdots & 0 \\ \vdots & \vdots & \vdots & \vdots & \ddots & \vdots \\ 0 & 0 & 0 & 0 & \cdots & 0 \end{pmatrix},
$$

$$
V_{EL} = \begin{pmatrix} 0 & 0 & 0 & 0 & \cdots & 0 \\ 0 & 0 & -q_1(1 - A_1^*/K_A)e^{-A_1^*/K_A}S_ES_L & -(q_2 - q_1A_1^*/K_A)e^{-A_1^*/K_A}S_ES_L & \cdots & (q_n - q_1A_1^*/K_A)e^{-A_1^*/K_A}S_ES_L \\ 0 & 0 & 0 & 0 & \cdots & 0 \\ 0 & 0 & 0 & 0 & \cdots & 0 \\ \vdots & \vdots & \vdots & \vdots & \ddots & \vdots \\ 0 & 0 & 0 & 0 & \cdots & 0 \end{pmatrix},
$$

$$
V_{ELPJ} = \begin{pmatrix} 0 & 0 & 0 & 0 & \cdots & 0 \\ 0 & 0 & 0 & 0 & \cdots & 0 \\ 0 & 0 & q_1\left(1 - \frac{A_1^*}{K_A}\right)e^{-A_1^*/K_A}S_ES_LS_{P1}S_J & \left(q_2 - q_1\frac{A_1^*}{K_A}\right)e^{-A_1^*/K_A}S_ES_LS_{P1}S_J & \cdots & \left(q_n - q_1\frac{A_1^*}{K_A}\right)e^{-A_1^*/K_A}S_ES_LS_{P1}S_J \\ 0 & 0 & 0 & 0 & \cdots & 0 \\ \vdots & \vdots & \vdots & \vdots & \ddots & \vdots \\ 0 & 0 & 0 & 0 & \cdots & 0 \end{pmatrix}.
$$

The matrices are upper triangular and therefore the characteristic equation for Equations S.2.15 simplifies to:

$$
\lambda(\lambda + \delta_L)(\lambda + \delta_A)^{n-1}\left(\lambda + \delta_A - \delta_A\left(1 - \ln\left(\frac{q_1S_ES_LS_{P1}S_J}{\delta_A}\right)\right)e^{-\tau\lambda}\right) = 0. \tag{S.2.17}
$$

The solutions to Equation (S.2.17) are $\lambda = 0$, $\lambda = -\delta_L$, $\lambda = -\delta_A$ and the solutions of

$$\lambda = -\delta_A + \delta_A \left(1 - \ln\left(\frac{q_1 S_E S_L S_{P1} S_J}{\delta_A}\right)\right) e^{-\tau\lambda}. \tag{S.2.18}$$

The stability of the persistence steady state is determined by Equation (S.2.18). As with the analysis of the extinction steady state, we apply Theorem 4.7 of Smith (2011) and conclude that the persistence steady state is unstable when $q_1 S_E S_L S_{P1} S_J < \delta_A$, this also corresponds to when the persistence state is biologically infeasible. When the persistence state is biologically feasible (i.e. $q_1 S_E S_L S_{P1} S_J > \delta_A$) then the steady state is stable provided $\ln(q_1 S_E S_L S_{P1} S_J/\delta_A) \leq 2$. When $\ln(q_1 S_E S_L S_{P1} S_J/\delta_A) \geq 2$ the stability of the persistence state depends on the length of the developmental delays.

In the case where $q_1 S_E S_L S_{P1} S_J > \delta_A$ and $\ln(q_1 S_E S_L S_{P1} S_J/\delta_A) \geq 2$ we consider the real solutions of Equation (S.2.18) and observe that there is either one of two real solutions and these are always negative; hence the stability of the persistence steady state must be lost by a pair of complex conjugate roots crossing the imaginary axis. We consider $\lambda = \mu i$, $\mu \in \mathbb{R}$ and equating real and imaginary parts, Equation (S.2.18) is reexpressed as:

$$(1 - \ln(q_1 S_E S_L S_{P1} S_J/\delta_A))\cos(\tau\mu) = 1, \tag{S.2.19a}$$

$$-\delta_A (1 - \ln(q_1 S_E S_L S_{P1} S_J/\delta_A))\sin(\tau\mu) = \mu. \tag{S.2.19b}$$

Rewriting these equations to eliminate $\mu$ gives:

$$2\pi k \pm \cos^{-1}\left(\frac{1}{1 - \ln(q_1 S_E S_L S_{P1} S_J/\delta_A)}\right) + \delta_A \tau \sqrt{(1 - \ln(q_1 S_E S_L S_{P1} S_J/\delta_A))^2 - 1} = 0, \tag{S.2.20}$$

where $k \in \mathbb{N}$. The stability boundary is determined by the first instance the eigenvalues cross the imaginary axis so we take $k = 0$. Therefore, the persistence steady state looses stability when:

$$-\cos^{-1}\left(\frac{1}{1 - \ln(q_1 S_E S_L S_{P1} S_J/\delta_a)}\right) + \delta_a \tau \sqrt{(1 - \ln(q_1 S_E S_L S_{P1} S_J/\delta_a))^2 - 1} < 0.$$

We summarise the results of the full stability analysis in Table S.2.2.

#### S.2.4 Close to size class boundaries

Figure S.2.1a illustrates the stability boundaries (Table S.2.2) in plastic trait space, namely in the maximum adult fecundity fecundity $q_1$ and through-pupal survival $S_{P1}$ parameter space. Treating $\alpha$ as a parameter we indicate how the traits vary with per capita larval food $\alpha$. We note that the low stress tipping point is not on a stability boundary, this is a consequence of an assumption made when conducting the linear stability analysis. We assumed $\alpha^*$ lies in the interior of a size-class and not at a boundary between classes. In practice, the assumption is violated when parameters are varied, however it is only violated when parameter changes result in a large change in $\alpha^*$ which occurs very close to a tipping point.

In Figure S.2.1b the continuous (in $\alpha$) form of the persistence steady state equation, where through-pupal survival and maximum adult fecundity are expressed as $S_P(\alpha^*)$ and $q(\alpha^*)$, is compared to the discrete size-class form of the steady state equations (obtained when $S_P(\alpha^*)$ and $q(\alpha^*)$ are replaced by $S_{Pk}$ and $q_k$). We would expect stability of the steady state to change at the turning point of the steady state curve, in practice the stability analysis predicts the change in stability occurs just past the turning point (Figure S.2.1c), this discrepancy is due to the violation of the assumption that $\alpha^*$ lies in the interior of a size-class and not at a boundary between size-classes. Of note, is that the discrepancy is small and the simplifying assumption provides an avenue for analysing a broader class of state and trait structured models.

| Steady state existence and stability | Conditions |
|---|---|
| (i) Extinction stable; Persistence not biologically feasible | $Q < D_A$ |
| (ii) Extinction unstable; Persistence stable | $Q > D_A$ and $\ln(Q/D_A) \leq 2$ |
| (iii) Extinction unstable; Persistence unstable node | $\ln(Q/D_A) \geq 2$ and $D_A\sqrt{(1-\ln(Q/D_A))^2 - 1} < \cos^{-1}\left(\frac{1}{1-\ln(Q/D_A)}\right)$ |
| (iv) Extinction unstable; Persistence unstable focus | $\ln(Q/D_A) \geq 2$ and $D_A\sqrt{(1-\ln(Q/D_A))^2 - 1} > \cos^{-1}\left(\frac{1}{1-\ln(Q/D_A)}\right)$ |

Table S.2.2: **Steady state existence and stability criteria**. We define $D_A = \delta_A(\tau_E + \tau_L + \tau_P + \tau_J)$ to be the adult stage mortality rate multiplied by the egg-adult development time and $Q = q_1 S_E S_L S_{J1}(\tau_E + \tau_L + \tau_P + \tau_J)$ to be the recruitment rate into the adult stage multiplied by the egg-adult development time.

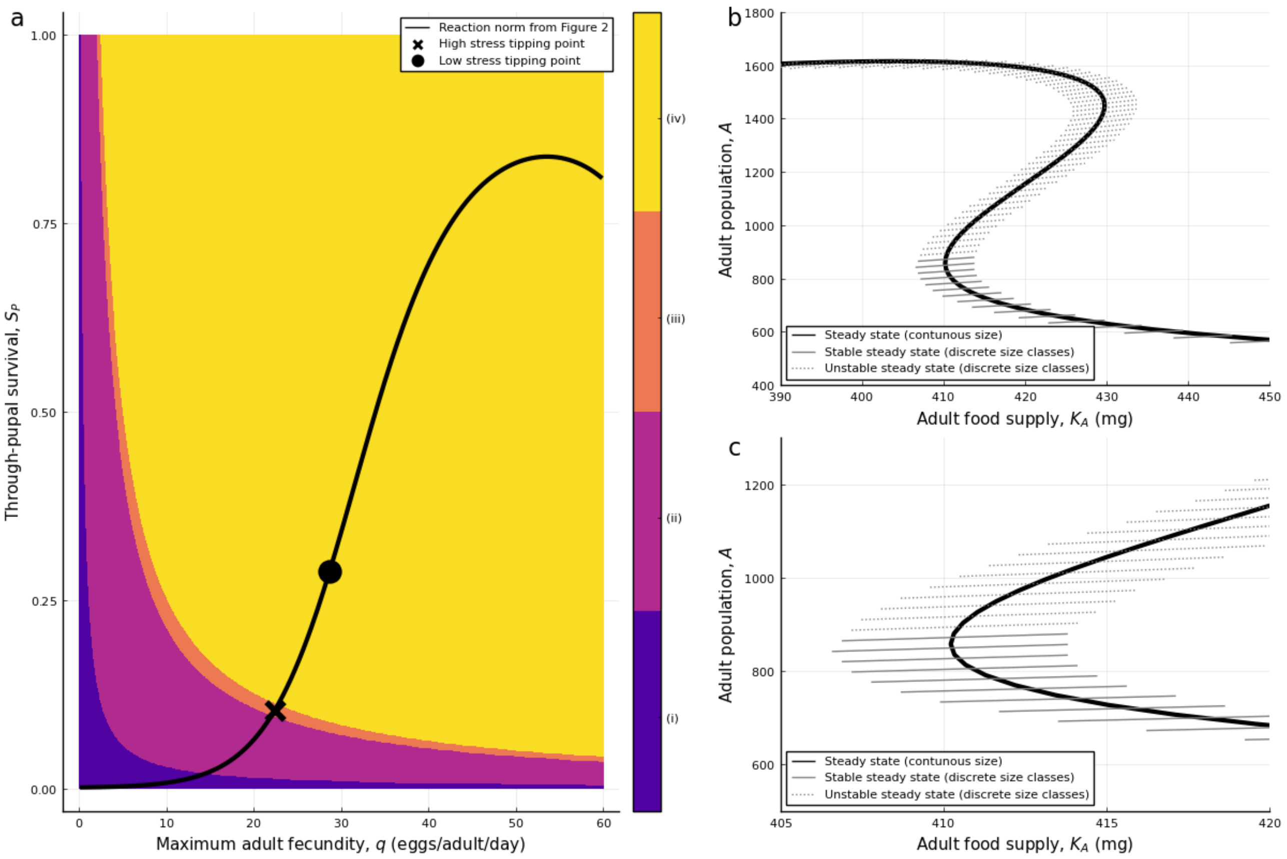


Figure S.2.1: **Comparison of analytical stability results to numerical simulations**. **a** The stability regions (as described in Table S.2.2) as through-juvenile survival $S_J$ and maximum adult fecundity $q$ are varied. The black line is the parametric solution to the reaction norms given in Equations (6) and (7) as $\alpha$ (the average food consumed through the larval stage) is varied, with $v_1 = -4$, $v_2 = 3.36$, and $v_3 = -0.50$. The high and low stress tipping points are marked with $\times$ and $\bullet$ respectively. **b** Adult population as the extrinsic environmental stress, adult food $K_A$, is slowly decreased and increased, demonstrating hysteresis. The solid thick line indicates the adult persistence steady state obtained from the continuous (in $\alpha$) form of the persistence steady state

### S.3 Parameter sensitivity analysis

The existence of tipping points is highly dependent on reaction norm shape and to explore this dependence further, we conducted a sensitivity analysis to study the sensitivity of tipping point existence to the reaction norm parameters from Equation (6) and Equation (7). A sample of 10,000 parameter sets was randomly chosen in the range of ±50% from the baseline values using Latin hypercube sampling (using the LatinHypercubeSampling package in Julia). For each parameter set in the sample we determined if model exhibits hysteresis (1) or not (0). Logistic regression was then performed on the categorical data using the GLM package in Julia, to determine which reaction norm parameters are the strongest predictors of the existence of tipping points (Table S.3.3). The sign of the coefficients from the logistic regression indicates whether increasing a parameter increases the likelihood of hysteresis, and the magnitude of the coefficient indicates the magnitude of the parameter effect. The $p$-value gives the likelihood that there is no relationship between the parameter and the model exhibiting tipping points, while a large $t$-value indicates the parameter is a predictor of tipping points, with positive values indicating an increased likelihood of tipping points. The only parameter that is not significant is $h_3$, whereas the existence of tipping points is most sensitive to $h_4$ and $v_3$. These parameters are responsible for the higher order non-linearity of the reaction norms.

| Reaction norm | | Coefficient | Standard Error | $t$-value | $p$-value |
|---|---|---|---|---|---|
| | (Intercept) | -7.48785 | 0.226261 | -33.09 | $< 1e{-99}$ |
| Maximum adult fecundity | $h_1$ | 0.252304 | 0.0173787 | 14.52 | $< 1e{-47}$ |
| | $h_2$ | 0.125606 | 0.010178 | 12.34 | $< 1e{-34}$ |
| | $h_3$ | 0.0252172 | 0.0696266 | 0.36 | 0.7172 |
| | $h_4$ | 2.12061 | 0.0939651 | 22.57 | $< 1e{-99}$ |
| Through-pupal survival | $v_1$ | 0.315043 | 0.0178327 | 17.67 | $< 1e{-69}$ |
| | $v_2$ | -0.151793 | 0.0203091 | -7.47 | $< 1e{-13}$ |
| | $v_3$ | -9.18834 | 0.234264 | -39.22 | $< 1e{-99}$ |

Table S.3.3: **Logistic regression to determine the sensitivity of tipping point existence to changes in reaction norm parameters.**

To understand how the value of parameters and not just the change affects the existence of tipping points, a kernel density estimation (KDE) is used to visualise the distribution of parameters that give rise to tipping points (Figure S.3.2). Figure S.3.2c shows that $v_3$ is highly skewed, implying that a relatively higher value of $v_3$ is important for the induction of tipping points. We can also see that $v_2$ and $q_4$ (Figure S.3.2b,g) are also skewed and therefore important in the induction of tipping points. The skewness of these parameters in the violin plots aligns with the results of the logistic regression presented in Table S.3.3.

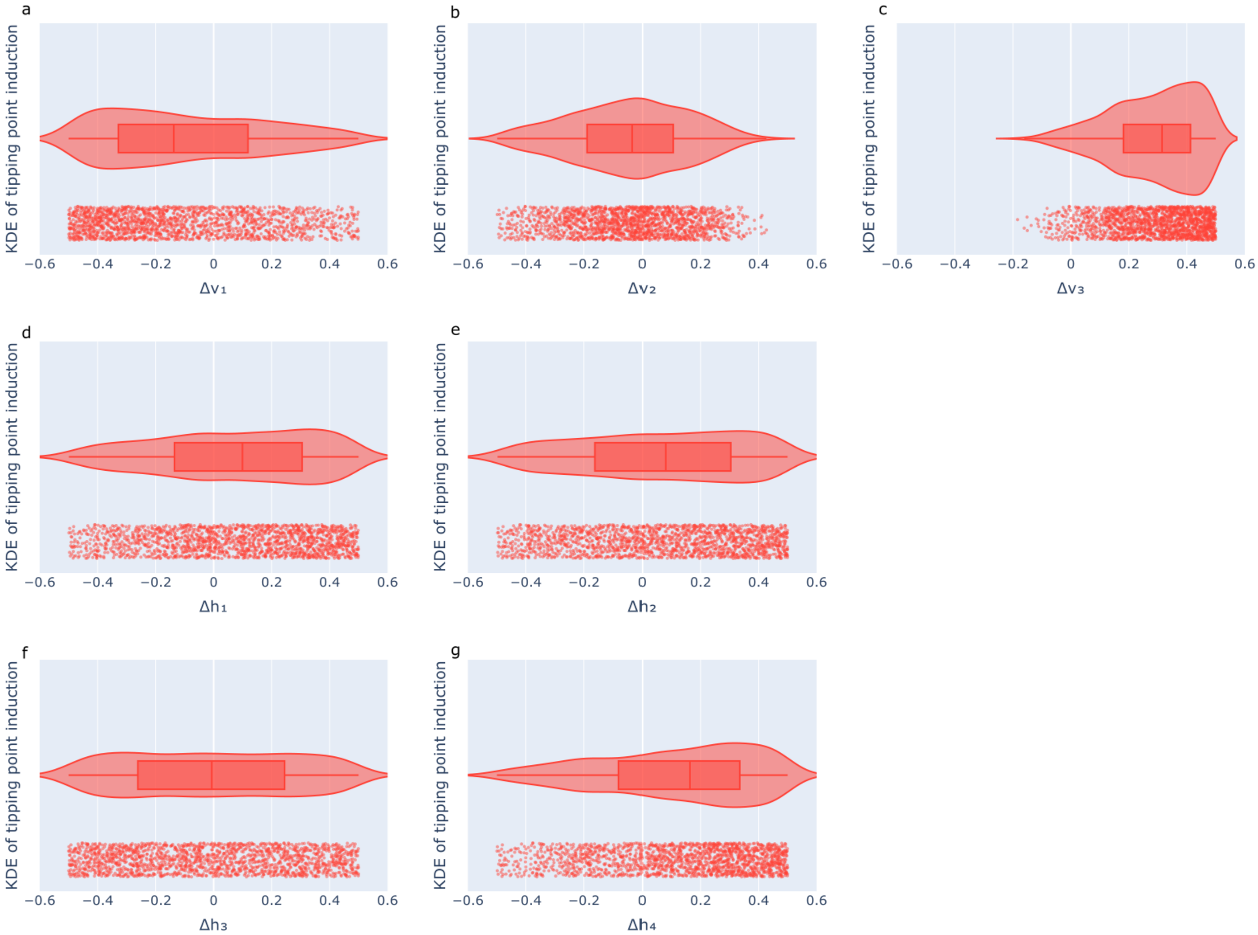


Figure S.3.2: **Sensitivity analysis of tipping point existence to parameters in the reaction norms, Equations (6) and (7).** Violin plots show the kernel density estimation (KDE) for each reaction norm parameter. Below each violin plot, the parameter set that induces tipping points is plotted. Parameters were chosen between ±50% from the baseline value using the methodology described in Table S.3.3. **a-c** parameters from the through-juvenile survival reaction norm, **d-g** parameters from the maximum adult fecundity reaction norm. Non-uniformity and skewness indicate the relative importance of a particular parameter to the induction of tipping points, as seen in **c** but not **e**.

#### S.3.1 Two parameter variation

The existence of tipping points, depending on parameter pairs was considered for each pair of reaction norm parameters, varying the parameters by ±50% from their baseline value, while keeping all the other parameters fixed. As the base parameter set does not result in tipping points, not all parameter pair variations result in the onset of tipping point. The parameter pair variations that did result in tipping points are presented in Figures S.3.3 to S.3.14. The only pairing of maximum adult fecundity parameters that induced tipping points was $h_2$ paired with $h_4$ (Figure S.3.4), which produced a narrow range of parameter space in which tipping points were observed. All other parameter pairs that gave rise to tipping points involved at least one through-pupal survival parameter.

To relate the location of tipping points to the underlying shapes of the reaction norms, a slice of two-dimensional parameter space was taken and the corresponding reaction norms were plotted (subplots b-c in Figures S.3.3 to S.3.14), together with the location of the low and high stress tipping points when these existed. In Figure S.3.4b-c, where only maximum adult fecundity parameters are varied, we observe that a sharp increase in the maximum adult fecundity separates the low and high stress tipping points. For all other parameter pair variations that led to tipping points, through-pupal survival parameters were included in the pair (Figures S.3.3 and S.3.5 to S.3.14), and we observe that through-pupal survival is suppressed at the tipping points.

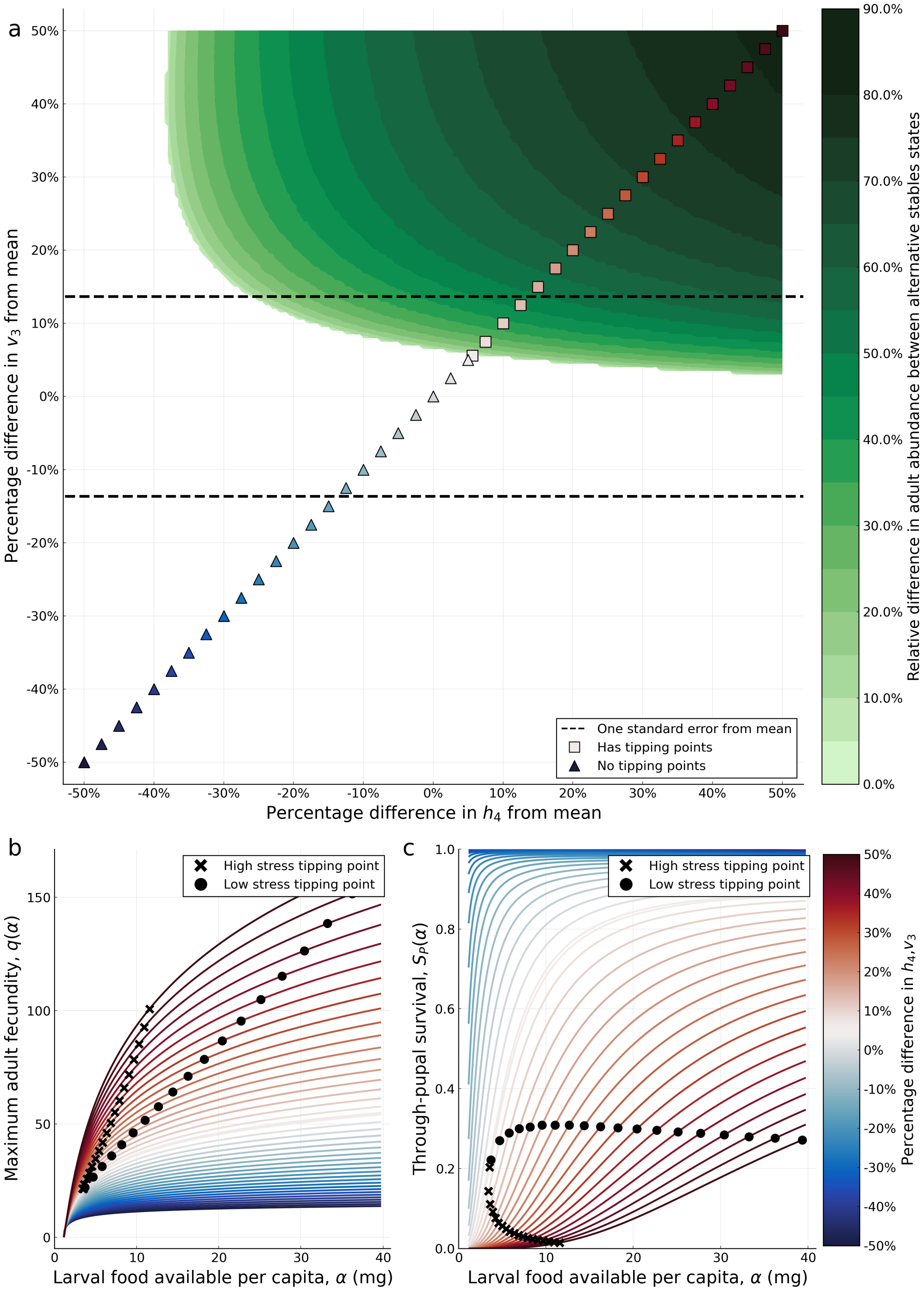


Figure S.3.3: **Effect of varying reaction norm parameters $h_4$ and $v_3$ on the existence of tipping points.** The parameters $h_4$ and $v_3$ control higher order non-linearity in the maximum adult fecundity and survival reaction norms respectively. (**a**), regions of $h_4$-$v_3$ parameter space in which tipping points occur are indicated in green, with the shading reflecting the relative change in adult abundance between the two tipping points. Dark shading indicates strong hysteresis. The reaction norm parameters $h_4$ and $v_3$ are varied by ±50% from their baseline values and all other parameters are held at their baseline values. The dashed line indicates one standard error from the estimated mean of $v_3$ (Table S.1.1). No data for standard errors for $h_4$ are available. (**b**)-(**c**), maximum adult fecundity ($q(\alpha)$) and through-pupal survival reaction norms ($S_P(\alpha)$) corresponding to the slice of two-dimensional parameter space indicated by the symbols in (**a**), where the parameter slice intersects with the tipping point existence region the location of the low and high stress tipping points are denoted crosses and dots respectively.

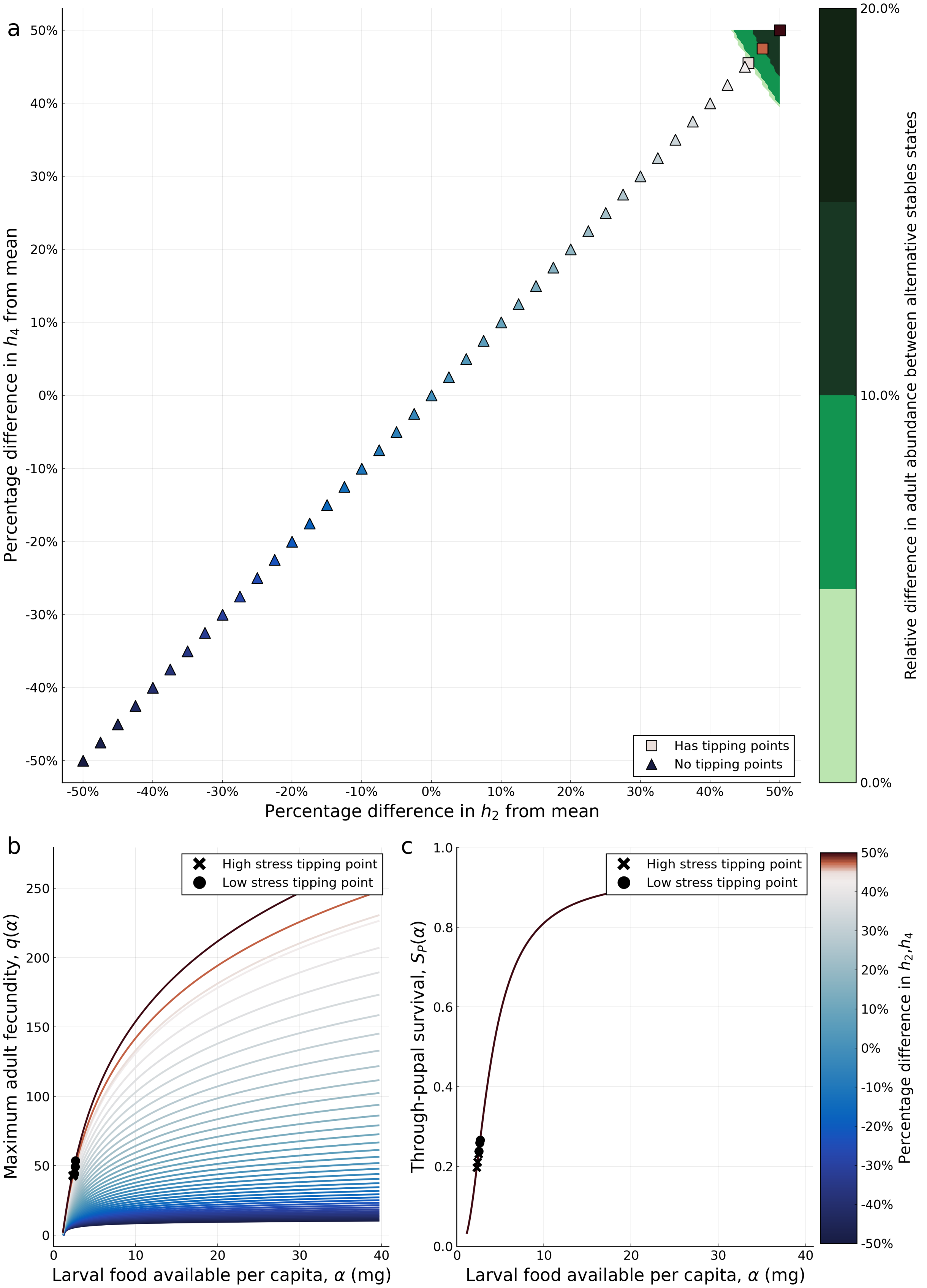


Figure S.3.4: **Effect of varying $h_2$ and $h_4$ on the existence of tipping points.** **a**, regions of $h_2$-$h_4$ parameter space in which tipping points occur are indicated in green, with the shading reflecting the relative change in adult abundance between the two tipping points. The maximum adult fecundity reaction norm parameters $h_2$ and $h_4$ are varied by $\pm 50\%$ from their baseline values and all other parameters are held at their baseline values. **b-c**, maximum adult fecundity and through-juvenile survival reaction norms corresponding to the slice of two-dimensional parameter space indicated in **a**, where the slice intersected with the tipping point existence region the location of the low and high stress tipping points are denoted by a cross and dot respectively.

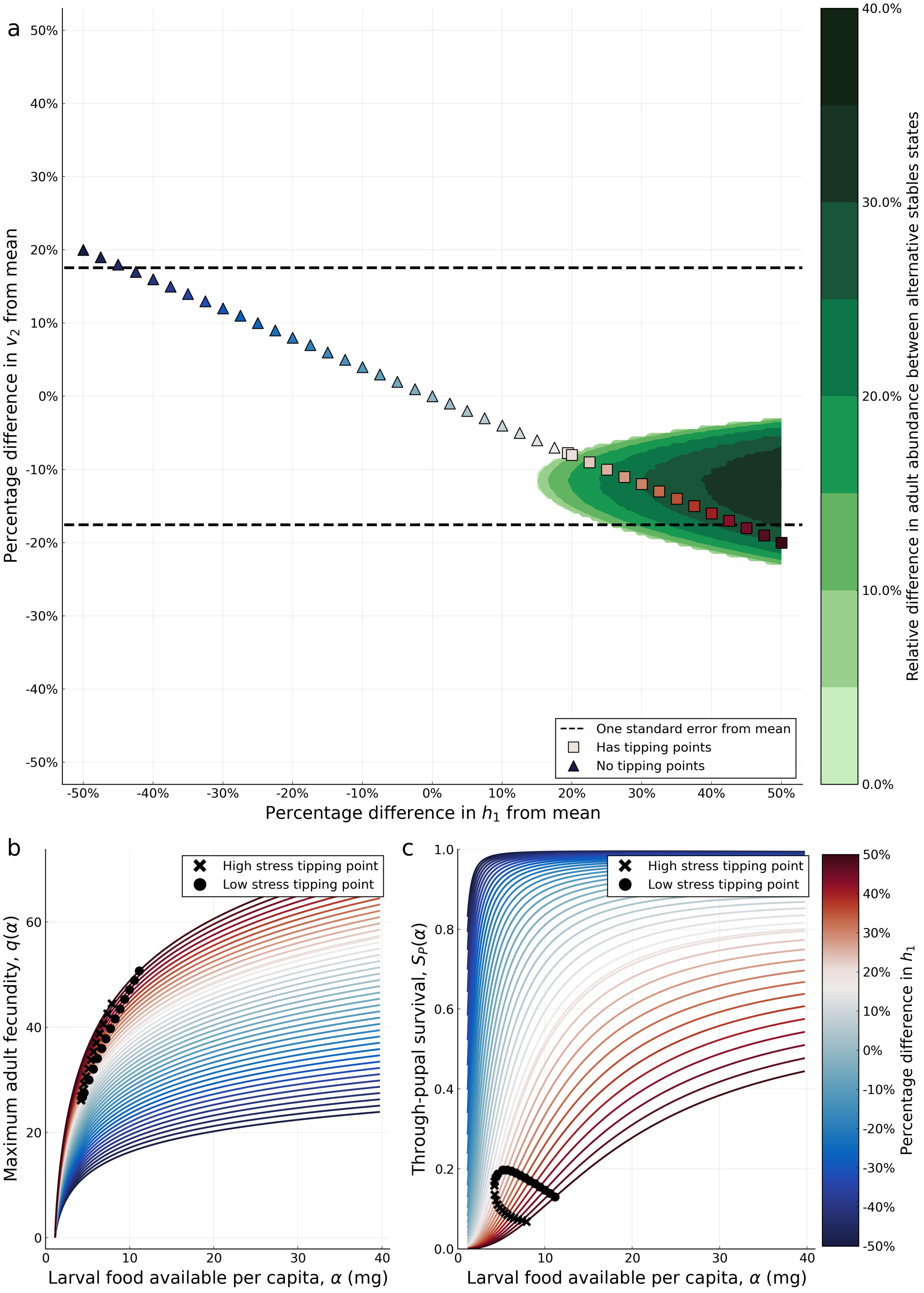


Figure S.3.5: **Effect of varying $h_1$ and $v_2$ on the existence of tipping points.** **a**, regions of $h_1$-$v_2$ parameter space in which tipping points occur are indicated in green, with the shading reflecting the relative change in adult abundance between the two tipping points. The parameters $h_1$ and $v_2$ are varied by $\pm 50\%$ from their baseline values and all other parameters are held at their baseline values. **b-c**, maximum adult fecundity and through-juvenile survival reaction norms corresponding to the slice of two-dimensional parameter space indicated in **a**, where the slice intersected with the tipping point existence region the location of the low and high stress tipping points are denoted by a cross and dot respectively.

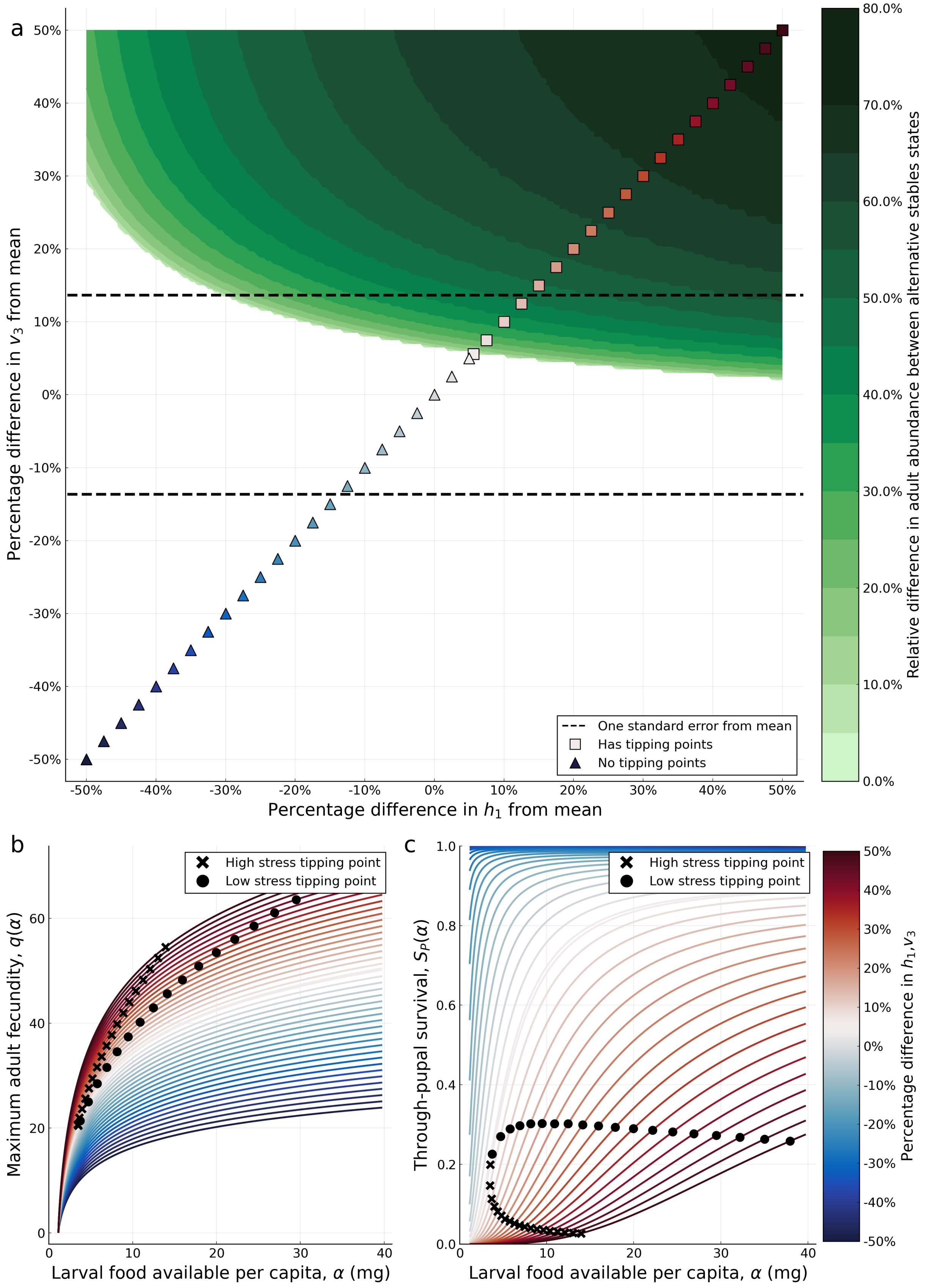


Figure S.3.6: **Effect of varying $h_1$ and $v_3$ on the existence of tipping points.** **a**, regions of $h_1$-$v_3$ parameter space in which tipping points occur are indicated in green, with the shading reflecting the relative change in adult abundance between the two tipping points. The parameters $h_1$ and $v_3$ are varied by $\pm 50\%$ from their baseline values and all other parameters are held at their baseline values. **b-c**, maximum adult fecundity and through-juvenile survival reaction norms corresponding to the slice of two-dimensional parameter space indicated in **a**, where the slice intersected with the tipping point existence region the location of the low and high stress tipping points are denoted by a cross and dot respectively.

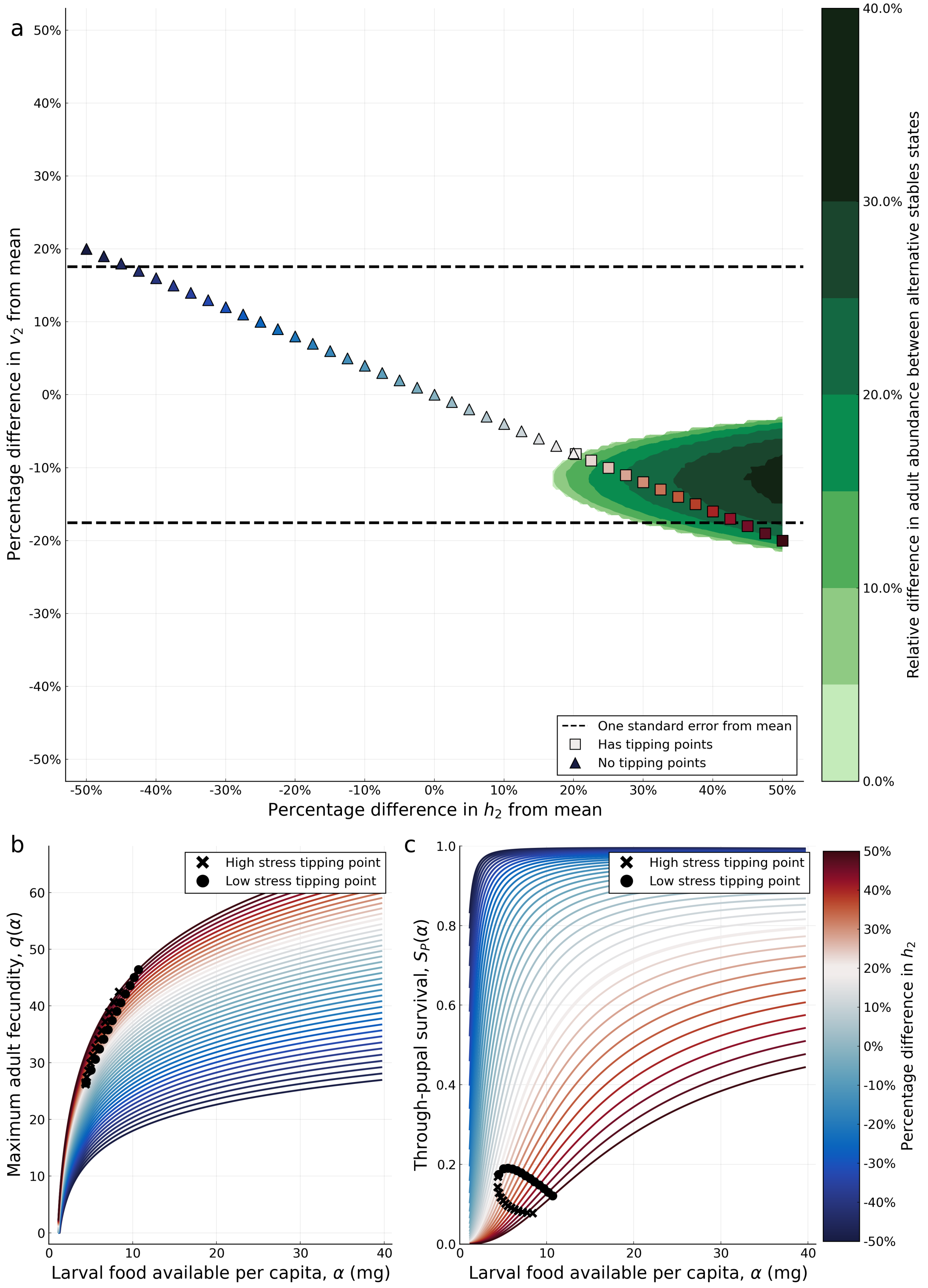


Figure S.3.7: **Effect of varying $h_2$ and $v_2$ on the existence of tipping points.** **a**, regions of $h_2$-$v_2$ parameter space in which tipping points occur are indicated in green, with the shading reflecting the relative change in adult abundance between the two tipping points. The parameters $h_2$ and $v_2$ are varied by $\pm 50\%$ from their baseline values and all other parameters are held at their baseline values. **b-c**, maximum adult fecundity and through-juvenile survival reaction norms corresponding to the slice of two-dimensional parameter space indicated in **a**, where the slice intersected with the tipping point existence region the location of the low and high stress tipping points are denoted by a cross and dot respectively.

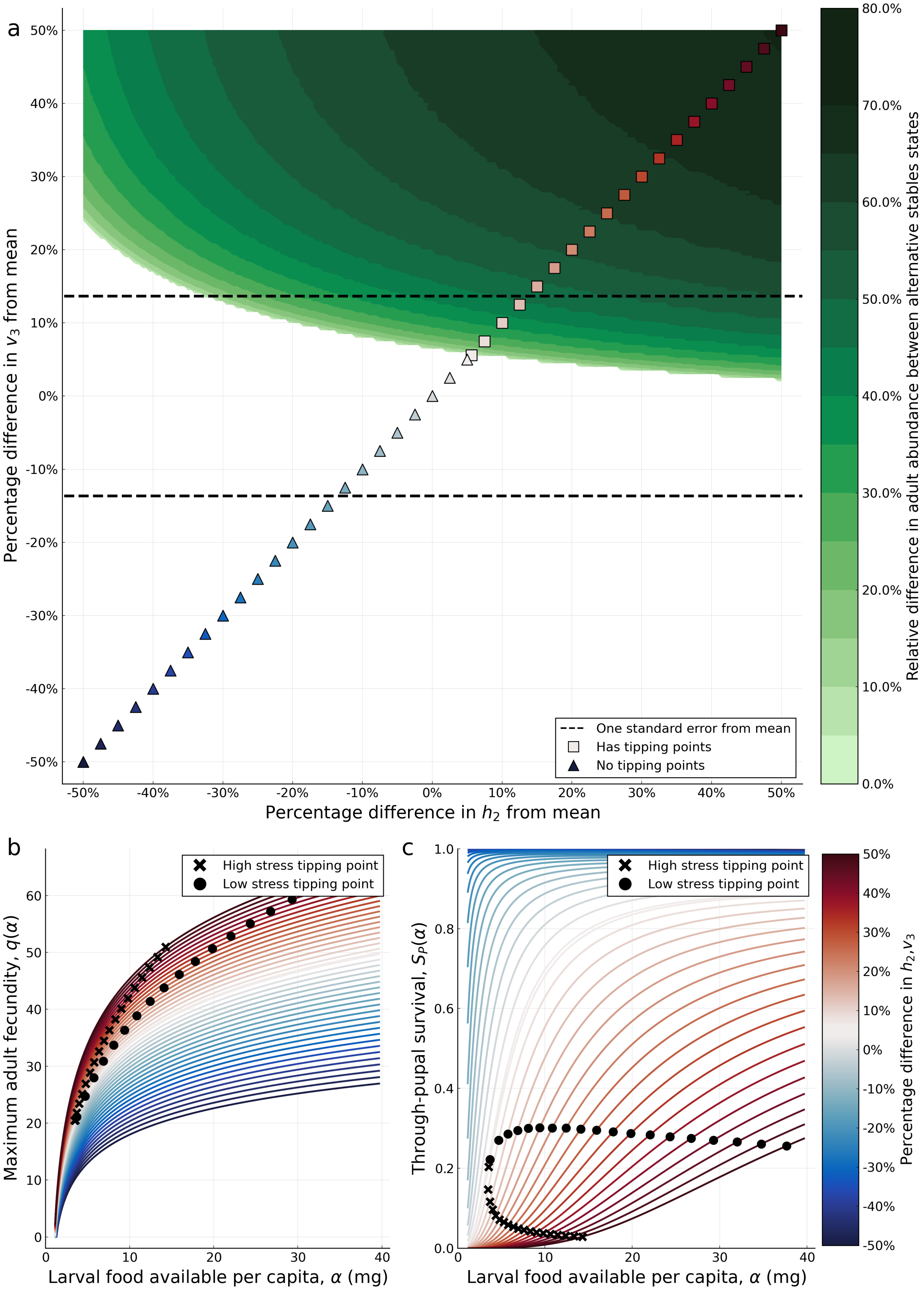


Figure S.3.8: **Effect of varying $h_2$ and $v_3$ on the existence of tipping points.** **a**, regions of $h_2$-$v_3$ parameter space in which tipping points occur are indicated in green, with the shading reflecting the relative change in adult abundance between the two tipping points. The parameters $h_3$ and $v_3$ are varied by $\pm 50\%$ from their baseline values and all other parameters are held at their baseline values. **b-c**, maximum adult fecundity and through-juvenile survival reaction norms corresponding to the slice of two-dimensional parameter space indicated in **a**, where the slice intersected with the tipping point existence region the location of the low and high stress tipping points are denoted by a cross and dot respectively.

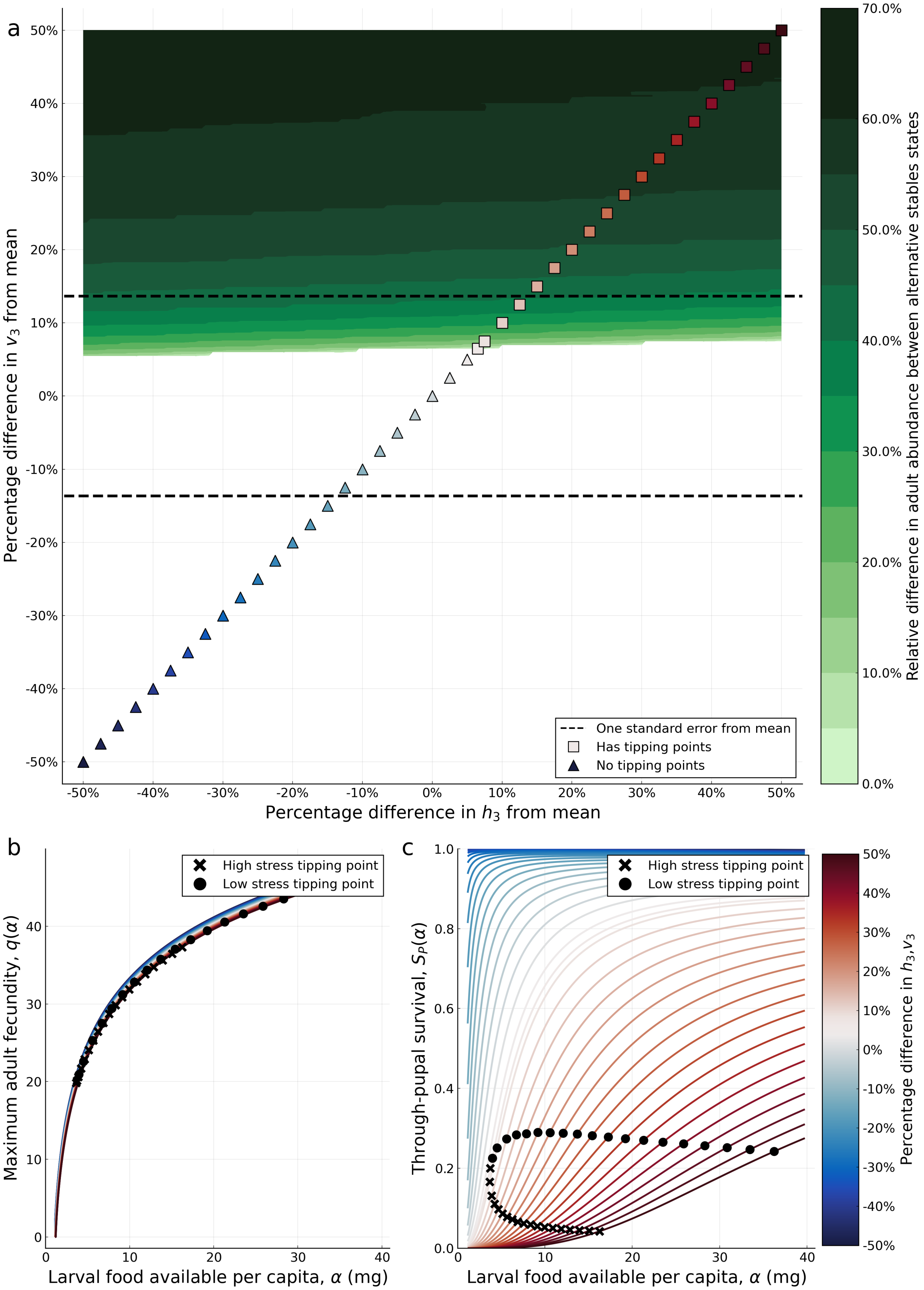


Figure S.3.9: **Effect of varying $h_3$ and $v_3$ on the existence of tipping points.** **a**, regions of $h_3$-$v_3$ parameter space in which tipping points occur are indicated in green, with the shading reflecting the relative change in adult abundance between the two tipping points. The parameters $h_3$ and $v_3$ are varied by $\pm 50\%$ from their baseline values and all other parameters are held at their baseline values. **b-c**, maximum adult fecundity and through-juvenile survival reaction norms corresponding to the slice of two-dimensional parameter space indicated in **a**, where the slice intersected with the tipping point existence region the location of the low and high stress tipping points are denoted by a cross and dot respectively.

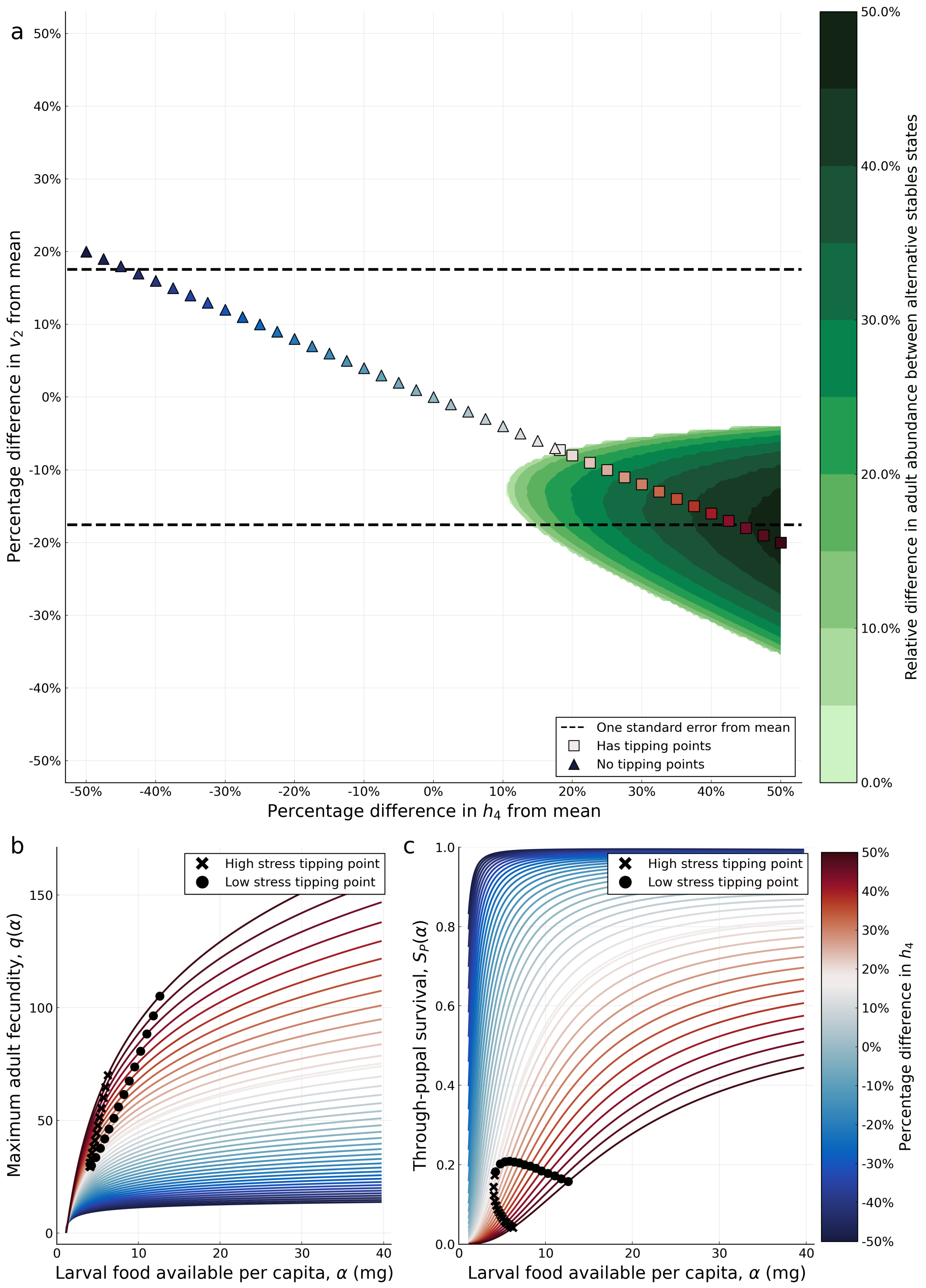


Figure S.3.10: **Effect of varying $h_4$ and $v_2$ on the existence of tipping points.** **a**, regions of $h_4$-$v_2$ parameter space in which tipping points occur are indicated in green, with the shading reflecting the relative change in adult abundance between the two tipping points. The parameters $h_4$ and $v_2$ are varied by ±50% from their baseline values and all other parameters are held at their baseline values. **b-c**, maximum adult fecundity and through-juvenile survival reaction norms corresponding to the slice of two-dimensional parameter space indicated in **a**, where the slice intersected with the tipping point existence region the location of the low and high stress tipping points are denoted by a cross and dot respectively.

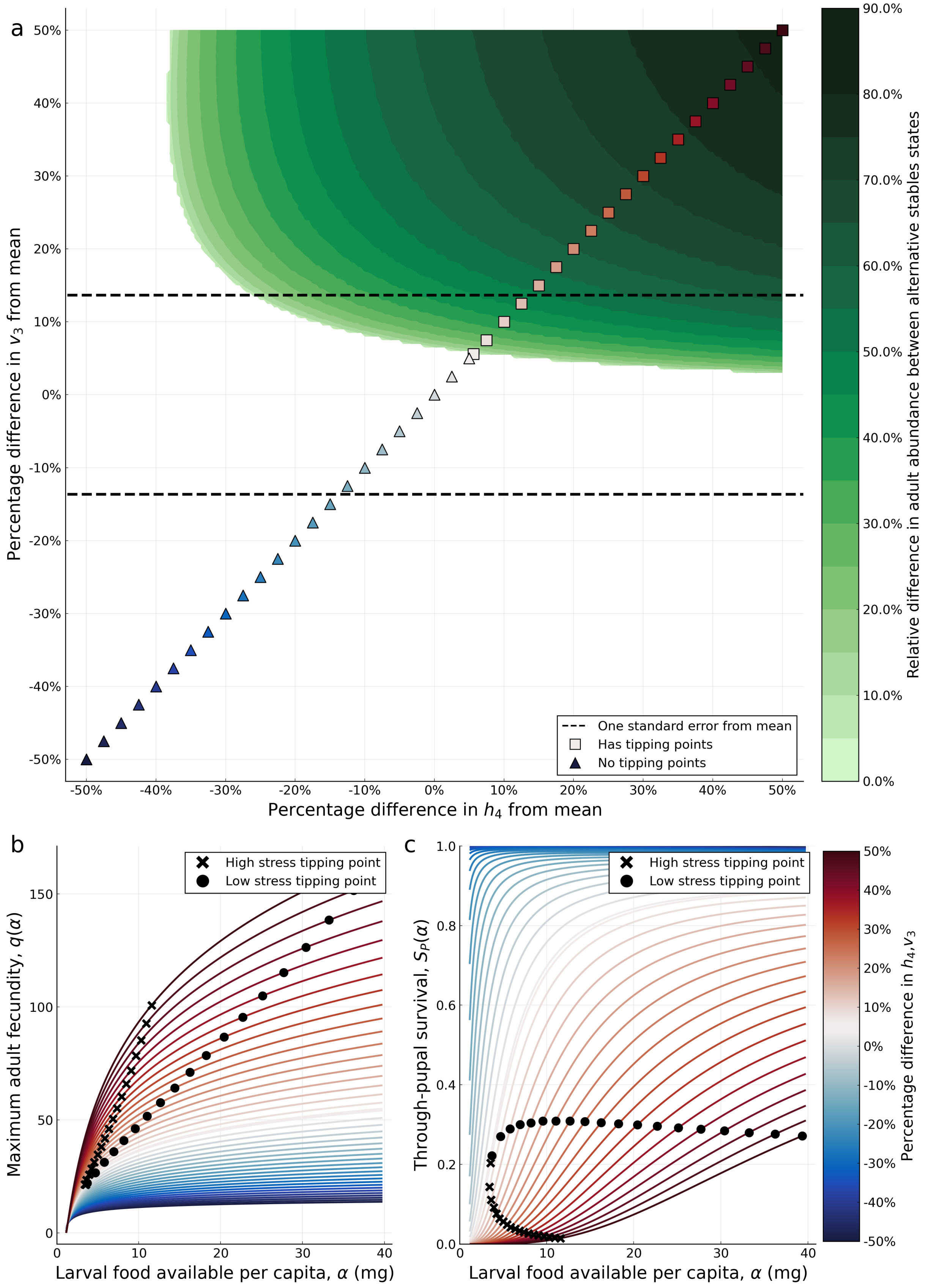


Figure S.3.11: **Effect of varying $h_4$ and $v_3$ on the existence of tipping points.** **a**, regions of $h_4$-$v_3$ parameter space in which tipping points occur are indicated in green, with the shading reflecting the relative change in adult abundance between the two tipping points. The parameters $h_4$ and $v_3$ are varied by ±50% from their baseline values and all other parameters are held at their baseline values. **b-c**, maximum adult fecundity and through-juvenile survival reaction norms corresponding to the slice of two-dimensional parameter space indicated in **a**, where the slice intersected with the tipping point existence region the location of the low and high stress tipping points are denoted by a cross and dot respectively.

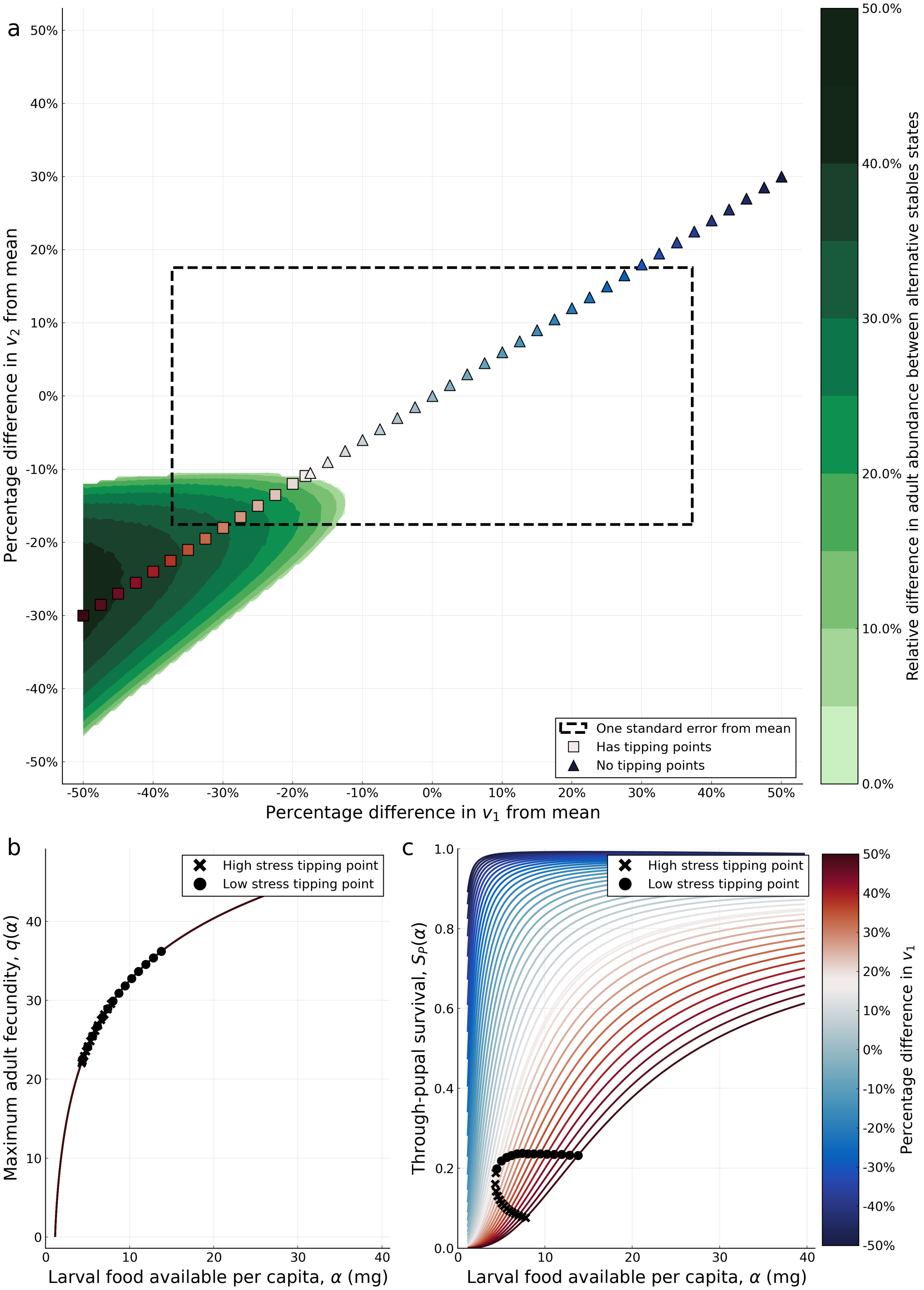


Figure S.3.12: **Effect of varying $v_1$ and $v_2$ on the existence of tipping points.** **a**, regions of $c_1$-$v_2$ parameter space in which tipping points occur are indicated in green, with the shading reflecting the relative change in adult abundance between the two tipping points. The parameters $v_1$ and $v_2$ are varied by ±50% from their baseline values and all other parameters are held at their baseline values. **b-c**, maximum adult fecundity and through-juvenile survival reaction norms corresponding to the slice of two-dimensional parameter space indicated in **a**, where the slice intersected with the tipping point existence region the location of the low and high stress tipping points are denoted by a cross and dot respectively.

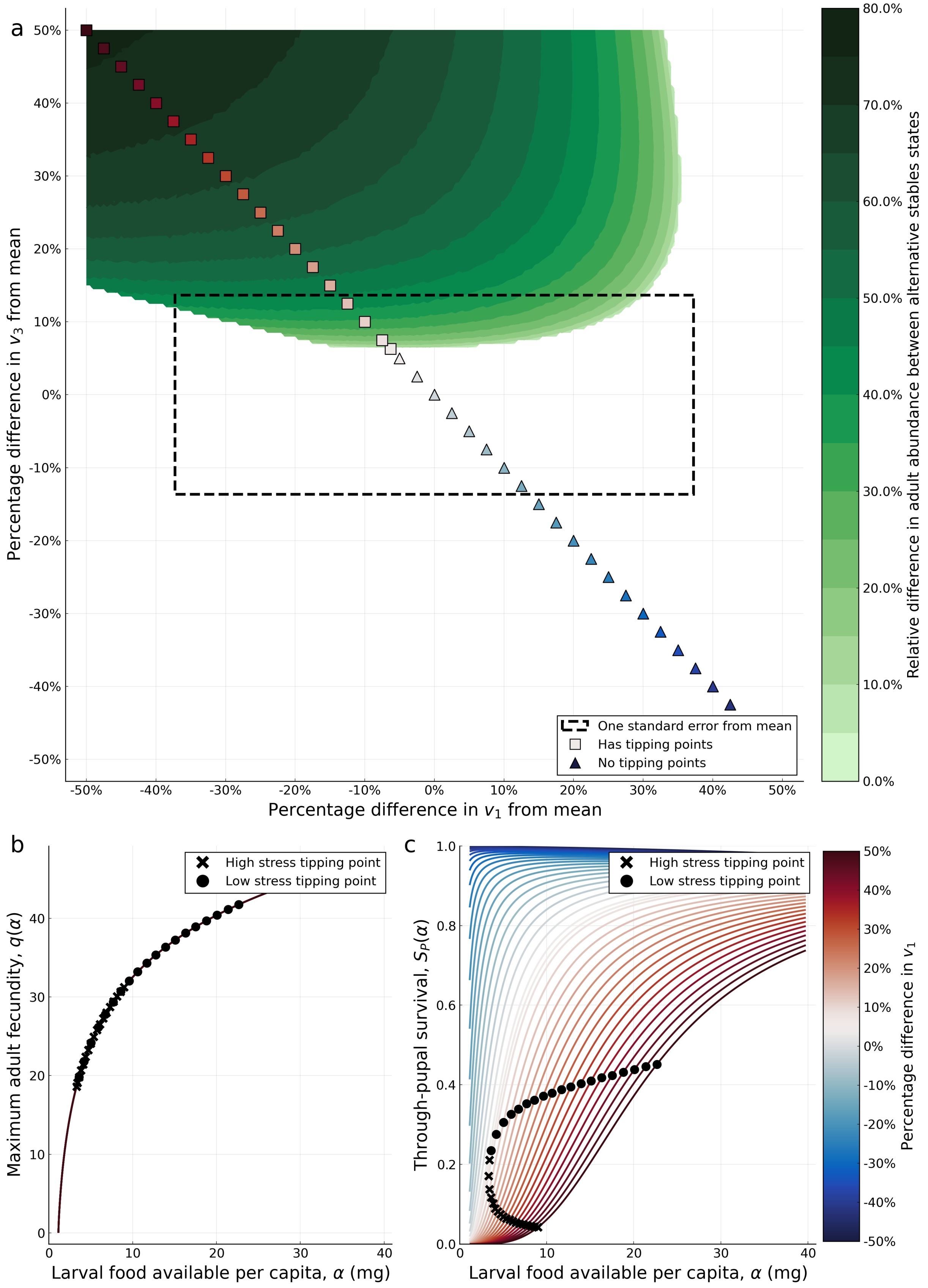


Figure S.3.13: **Effect of varying $v_1$ and $v_3$ on the existence of tipping points.** **a**, regions of $v_1$-$v_3$ parameter space in which tipping points occur are indicated in green, with the shading reflecting the relative change in adult abundance between the two tipping points. The parameters $v_1$ and $v_3$ are varied by ±50% from their baseline values and all other parameters are held at their baseline values. **b-c**, maximum adult fecundity and through-juvenile survival reaction norms corresponding to the slice of two-dimensional parameter space indicated in **a**, where the slice intersected with the tipping point existence region the location of the low and high stress tipping points are denoted by a cross and dot respectively.

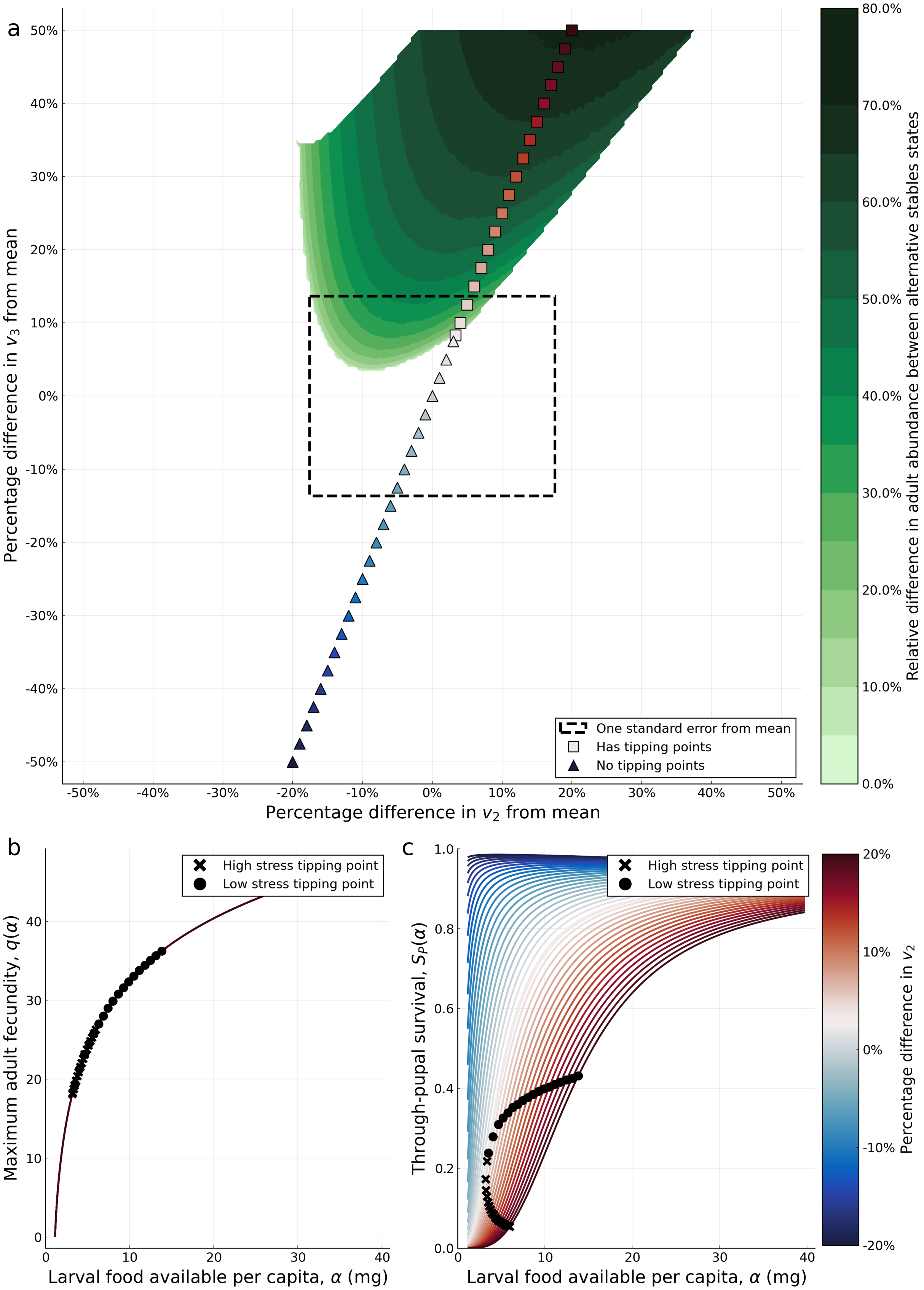


Figure S.3.14: **Effect of varying $v_2$ and $v_3$ on the existence of tipping points.** **a**, regions of $v_2$-$v_3$ parameter space in which tipping points occur are indicated in green, with the shading reflecting the relative change in adult abundance between the two tipping points. The parameters $v_2$ and $v_3$ are varied by $\pm 50\%$ from their baseline values and all other parameters are held at their baseline values. **b-c**, maximum adult fecundity and through-juvenile survival reaction norms corresponding to the slice of two-dimensional parameter space indicated in **a**, where the slice intersected with the tipping point existence region the location of the low and high stress tipping points are denoted by a cross and dot respectively.

### S.4 Shared resource model

An ontogenic niche shift does not occur if resources are shared between the larvae and adults. If we assume that adults and larvae compete for a shared food source that is supplied daily at rate $K$ then the equation describing the per captia larval food (Equation (3)) and the adult fecundity term in Equation (2) are modified account for the competition between the larvae and adults, with a constant $c$ that determines the relative ability of larvae and adults to compete for food.

To determine if hysteresis can occur in this shared resource model, we compute the persistence steady state for the updated model satisfied by:

$$0 = q_k S_E(1 - S_L)A^* \exp\left(-\frac{L^* + cA^*}{K}\right) - \delta_L L^*, \tag{S.4.21a}$$

$$0 = A_k^* \left(q_k S_E S_L S_{Pk} S_J \exp\left(-\frac{L^* + cA^*}{K}\right) - \delta_A\right), \tag{S.4.21b}$$

$$0 = A_i^* \text{ for } i = 1, ..., k-1, k+1, ..., n, \tag{S.4.21c}$$

where food per capita satisfies

$$\alpha^* = \frac{K}{L^* + cA^*}. \tag{S.4.22}$$

Expressing $q_k$ and $S_{Pk}$ in terms of their continuous functional equivalents, $q(\alpha)$ and $S_P(\alpha)$, allows Equation (S.4.21b) to be expressed in terms of $\alpha^*$ to give:

$$\frac{q(\alpha^*)S_E S_L S_P(\alpha^*)S_J}{\delta_A} = \exp\left(\frac{1}{\alpha^*}\right). \tag{S.4.23}$$

The solutions to Equation (S.4.23) give the values of $\alpha^*$, which is independent of $K$ and so when food supply, $K$ is changed, the larval and adult populations change proportionally.

Rearranging Equation (S.4.21a) and expressing $L^*$ in terms of $A$ and $\alpha^*$ we obtain an equation for the adult steady state:

$$A^* = \frac{\delta_L K}{\alpha^*\left(q(\alpha^*)S_E(1 - S_L)\exp\left(-\frac{1}{\alpha^*}\right) + c\delta_L\right)}. \tag{S.4.24}$$

and observe that $A^*$ changes linearly with $K$. Similarly, the linear dependence of $L^*$ on $K$ follows from rearranging Equation (S.4.22). Therefore, changes to food supply do not effect the number of persistence steady states and hysteresis cannot occur in the shared resource model.

### S.5 Trait sensitivity model

Two simplifying assumptions are made to the Brass *et al.* (2021) model, the first is to assume that fecundity is not plastic, so $q(\alpha)$ is a constant that we denote by $q$, second we assume the through-pupal survival reaction norm can be described by the Holling type 3 function:

$$S_P(\alpha) = \frac{a\alpha^2}{1 + ah\alpha^2}.$$

Under these simplifications the larval and adult nullclines are given by $dL/dt = 0$ and $dA/dt = 0$, which solve to give the non-trivial nullclines:

$$L = qS_E(1 - S_L)\exp(-A/K_A)A/\delta_L, \quad \text{Larval nullcline} \tag{S.5.25}$$

$$A = K_A \ln\left(\frac{qS_E S_L S_P(K_L/L)S_J}{\delta_A}\right), \quad \text{Adult nullcline.} \tag{S.5.26}$$

#### S.5.1 Properties of the adult nullcline

To understand the shape of the adult nullcline we first find the turning points of the adult nullcline and determine how the location of these points varies with $K_A$ and $K_L$. Differentiating Equation (S.5.26) with respect to $L$ gives:

$$\frac{\partial A}{\partial L} = -\frac{K_L K_A}{L^2}\frac{S_P'(K_L/L)}{S_P(K_L/L)}, \tag{S.5.27}$$

which is zero if and only if $S_P'(K_L/L) = 0$. The turning point condition fixes the value of the ratio $K_L/L$ and hence changing $K_L$ does not change the value of $A$ at which a turning point in the adult nullcline occurs, but does change the value of $L$ at which the turning point occurs. Changing $K_A$ linearly changes the value of $A$ at which the adult nullcline turning point occurs, but does not change the value of $L$ at which the turning point occurs.

The adult nullcline intersects the $A$-axis as $L \to 0$, approaching $A = K_A \ln\left(qS_E S_L S_P(\infty) S_J/\delta_A\right)$. Taking $S_P(\alpha)$ to be the Holling type-3 function yields $S_P(\infty) = 1/h$ adn then $A = K_A \ln\left(qS_E S_L S_J/h\delta_A\right)$. The adult nullcline intersects the $L$-axis when $A = 0$ giving $R_0(K_L/L) = qS_E S_L S_P(K_L/L)S_J/\delta_A = 1$, which is independent of $K_A$ and changes linearly with $K_L$. In summary, the adult nullcline is hump shaped function of $L$ if the reaction norm $S_P(\alpha)$ has turning point, but otherwise it is a decreasing function of $L$, intersecting the $A$-axis $A = K_A \ln(R_0(\infty))$, which depends on $K_A$, but not $K_l$ and intersects the $L$-axis at $R_0(\alpha) = 1$, which depends on $K_L$, bit not $K_A$.

#### S.5.2 Properties of the larval nullcline

To find turning points in the larval nullcline we differentiate Equation (S.5.25) by $A$:

$$\frac{\partial L}{\partial A} = \frac{L}{A}\left(1 - \frac{A}{K_A}\right), \tag{S.5.28}$$

which is equal to zero if and only if $L = 0$ or $A = K_A$. The larval nullcline, $L$ can be expressed as a function of $A$ that only has one root at the origin and on the larval nullcline $A = 0$ if and only if $L = 0$.

Considering what happens to the larval nullcline in the limit as $A \to \infty$ gives:

$$\lim_{A\to\infty}(L) = \lim_{A\to\infty}\left(qS_E(1 - S_L)\exp(-A/K_A)A/\delta_L\right) = 0. \tag{S.5.29}$$

In summary, the larval nullcline is hump shaped when viewed as function of $A$. The maximum occurs at $A = K_A$ and it increases with $K_A$ and the nullcline declines to $L = 0$ as $A \to \infty$. The larval nullcline does not change with $K_L$.

#### S.5.3 Boundary for hysteresis in $a$-$h$ parameter space

Hysteresis does not occur if of $h$ above $h_T$. We determine $h_T$ by noting that a prerequisite for hysteresis to occur is for Equation (S.2.2) to have at least one turning point, in other words we need $\partial K_A/\partial\alpha^* = 0$. The turning point is lost when $\partial^2 K_A/\partial\alpha^{*2} = 0$ and hence $h_T$ can be found by simultaneously solving $\partial K_A/\partial\alpha^* = 0$ and $\partial^2 K_A/\partial\alpha^{*2} = 0$. When $q$ is constant simplifies to:

$$\frac{\partial K_A}{\partial\alpha^*} = K_A\left(\frac{S_P'(\alpha^*)}{S_P(\alpha^*)} - \frac{1}{\alpha^*} - \frac{1}{\ln\left(\frac{q(\alpha^*)S_E S_L S_P(\alpha^*)S_J}{\delta_A}\right)}\left(\frac{S_P'(\alpha^*)}{S_P(\alpha^*)}\right)\right). \tag{S.5.30}$$

Taking the second derivative of $K_A$ with respect to $\alpha^*$, we obtain:

$$\begin{aligned}\frac{\partial^2 K_A}{\partial \alpha^{*2}} =& \frac{\partial K_A}{\partial \alpha^*}\left(\frac{S_P'(\alpha^*)}{S_P(\alpha^*)} - \frac{1}{\alpha^*} - \frac{1}{\ln\left(\frac{q(\alpha^*)S_E S_L S_P(\alpha^*)S_J}{\delta_A}\right)}\left(\frac{S_P'(\alpha^*)}{S_P(\alpha^*)}\right)\right) \\ &+ K_A\left(\frac{S_P''(\alpha^*)}{S_P(\alpha^*)} - \left(\frac{S_P'(\alpha^*)}{S_P(\alpha^*)}\right)^2 + \frac{1}{\alpha^2}\right. \\ &\left.+ \frac{1}{\ln\left(\frac{q(\alpha^*)S_E S_L S_P(\alpha^*)S_J}{\delta_A}\right)^2}\left(\frac{S_P'(\alpha^*)}{S_P(\alpha^*)}\right)^2 - \frac{1}{\ln\left(\frac{q(\alpha^*)S_E S_L S_P(\alpha^*)S_J}{\delta_A}\right)}\left(\frac{S_P''(\alpha^*)}{S_P(\alpha^*)} - \left(\frac{S_P'(\alpha^*)}{S_P(\alpha^*)}\right)^2\right)\right).\end{aligned} \tag{S.5.31}$$

Then if we take $dK_A/d\alpha^* = d^2K_A/d\alpha^{*2} = 0$, we obtain the equation:

$$0 = \left(\frac{S_P''(\alpha^*)}{S_P(\alpha^*)} - \left(\frac{S_P'(\alpha^*)}{S_P(\alpha^*)}\right)^2\right)\frac{1}{\alpha^*}\frac{S_P(\alpha^*)}{S_P'(\alpha^*)} + \frac{1}{\alpha^2} + \left(\frac{S_P'(\alpha^*)}{S_P(\alpha^*)} - \frac{1}{\alpha^*}\right)^2. \tag{S.5.32}$$

Applying the Holling type-3 function for $S_P(\alpha)$, we can obtain $0 = 1 - 4ah\alpha^2 - a^2h^2\alpha^4$, solving this equation determines maximum value of $\alpha^*$ for which hysteresis can occur, $\alpha^* = \sqrt{((\sqrt{5}-2)/(ah))}$. Which can be substituted back into the hysteresis condition Equation (S.5.30) to obtain a threshold value for $h$:

$$h_T = \frac{3-\sqrt{5}}{4}\frac{qS_ES_LS_J}{\delta_A}e^{-\frac{3-\sqrt{5}}{2}} \approx 1.44 \text{ (3s.f.)} \tag{S.5.33}$$

Hysteresis occurs when $h < h_T$.

We can also calculate a upper bound on $S_P(\alpha)$ for existence of tipping points. The persistence steady state satisfies Equation (S.2.2) and tipping points are located where Equation (S.5.30) is zero. Solving these two equations for $q(\alpha) = q$ and $S_P(\alpha)$ given by the Holling type 3 function gives:

$$K_A = \frac{K_L\delta_LS_LS_J}{\delta_A(1-S_L)}\left(\frac{a\alpha(1-ah\alpha^2)}{2(1+ah\alpha^2)}\right).$$

To ensure $K_A > 0$ we require $\alpha^2 < a/h$, or equivalently the equation for $S_P(\alpha)$ gives $S_P(\alpha) < 1/2h$ and hence tipping points can only be located in the lower half of the through-pupal survival reaction norm. Moreover, since increasing $h$ lowers the threshold value of $S_P$, large values of $h$ will not give rise to tipping points as demonstrated by the condition that $h < h_T$.
